# Cohesin loss eliminates all loop domains, leading to links among superenhancers and downregulation of nearby genes

**DOI:** 10.1101/139782

**Authors:** Suhas S. P. Rao, Su-Chen Huang, Brian Glenn St Hilair, Jesse M. Engreitz, Elizabeth M. Perez, Kyong-Rim Kieffer-Kwon, Adrian L. Sanborn, Sarah E. Johnstone, Ivan D. Bochkov, Xingfan Huang, Muhammad S. Shamim, Arina D. Omer, Bradley E. Bernstein, Rafael Casellas, Eric S. Lander, Erez Lieberman Aiden

## Abstract

The human genome folds to create thousands of intervals, called “contact domains,” that exhibit enhanced contact frequency within themselves. “Loop domains” form because of tethering between two loci - almost always bound by CTCF and cohesin – lying on the same chromosome. “Compartment domains” form when genomic intervals with similar histone marks co-segregate. Here, we explore the effects of degrading cohesin. All loop domains are eliminated, but neither compartment domains nor histone marks are affected. Loci in different compartments that had been in the same loop domain become more segregated. Loss of loop domains does not lead to widespread ectopic gene activation, but does affect a significant minority of active genes. In particular, cohesin loss causes superenhancers to co-localize, forming hundreds of links within and across chromosomes, and affecting the regulation of nearby genes. Cohesin restoration quickly reverses these effects, consistent with a model where loop extrusion is rapid.

## INTRODUCTION

Many studies have shown that the insulator protein CTCF and the ring-shaped cohesin complex colocalize on chromatin (Parelho et al., 2008; Rubio et al., 2008; Wendt et al., 2008) and lie at the anchors of loops (Heidari et al., 2014; Rao et al., 2014; Splinter et al., 2006; Tang et al., 2015) and the boundaries of contact domains (sometimes called “topologically constrained domains”, “topologically associated domains”, or “physical domains”) (Dixon et al., 2012; Lieberman-Aiden et al., 2009; Nora et al., 2012; Phillips-Cremins et al., 2013; Rao et al., 2014; Sexton et al., 2012). These findings suggest that these proteins play a role in regulating genome folding (Benabdallah and Bickmore, 2015; Dekker and Misteli, 2015; Lupiáñez et al., 2016; Merkenschlager and Nora, 2016; Phillips-Cremins and Corces, 2013; Uhlmann, 2016). Similarly, deletion of individual CTCF sites can interfere with loop and contact domain formation (Guo et al., 2015; Narendra et al., 2015; Sanborn et al., 2015; de Wit et al., 2015). However, low-resolution experiments examining genome-wide depletion of CTCF and cohesin have thus far observed only limited effects on chromosome architecture, reporting that compartments and contact domains still appear to be present (Seitan et al., 2013; Sofueva et al., 2013; Zuin et al., 2014). These results have made it difficult to ascertain the role of CTCF and cohesin in the regulation of genome topology at the chromosome scale.

Here, we examine the effects of cohesin loss on nuclear architecture, epigenetic state, and transcription. By generating maps of DNA-DNA contacts of much higher resolution, we are able to characterize the effects of global cohesin loss on nuclear architecture more clearly than in earlier studies. We demonstrate that there are two types of domains: loop domains, which disappear within an hour of cohesin degradation; and compartment domains, which do not. Cohesin restoration quickly rescues the loop domains (<1 hour). In the absence of cohesin, we observe that superenhancers tend to co-segregate, forming hundreds of links both within and across chromosomes. Surprisingly, the transcriptional effects of cohesin degradation are limited to a small number of genes, many of which lie in close proximity to superenhancers.

## RESULTS

### Rapid degradation of RAD21 using an auxin-inducible degron system

To study the effects of cohesin loss on genome folding and gene expression, we employed an auxin-inducible degron (AID) (Natsume et al., 2016; Nishimura et al., 2009) to destroy RAD21, a core component of the cohesin ring complex. In this system, constitutive expression of the auxin-activated ubiquitin ligase TIR1 leads, in the presence of auxin, to rapid ubiquitination and degradation of proteins tagged with an AID domain. We used this system in HCT-116, a human colorectal carcinoma epithelial cell line. This cell line had been previously modified by Natsume *et al.* (Natsume et al., 2016) so that both alleles of RAD21 were tagged with an AID domain and a fluorescent mClover (RAD21-mAID-mClover, “RAD21-mAC”) (Fig. 1A). We confirmed that RAD21-mAC was efficiently degraded after 6 hours of auxin treatment using fluorescence microscopy and ChIP with antibodies for RAD21 (Fig. 1B, S1, see Methods). We also confirmed that the loss of RAD21 disrupted the ability of cohesin to associate with DNA by performing ChIP-Seq using antibodies for SMC1, a different subunit of the cohesin ring, and confirming the disappearance of SMC1 peaks across the genome (Fig. 1C, D).

**Figure 1:**
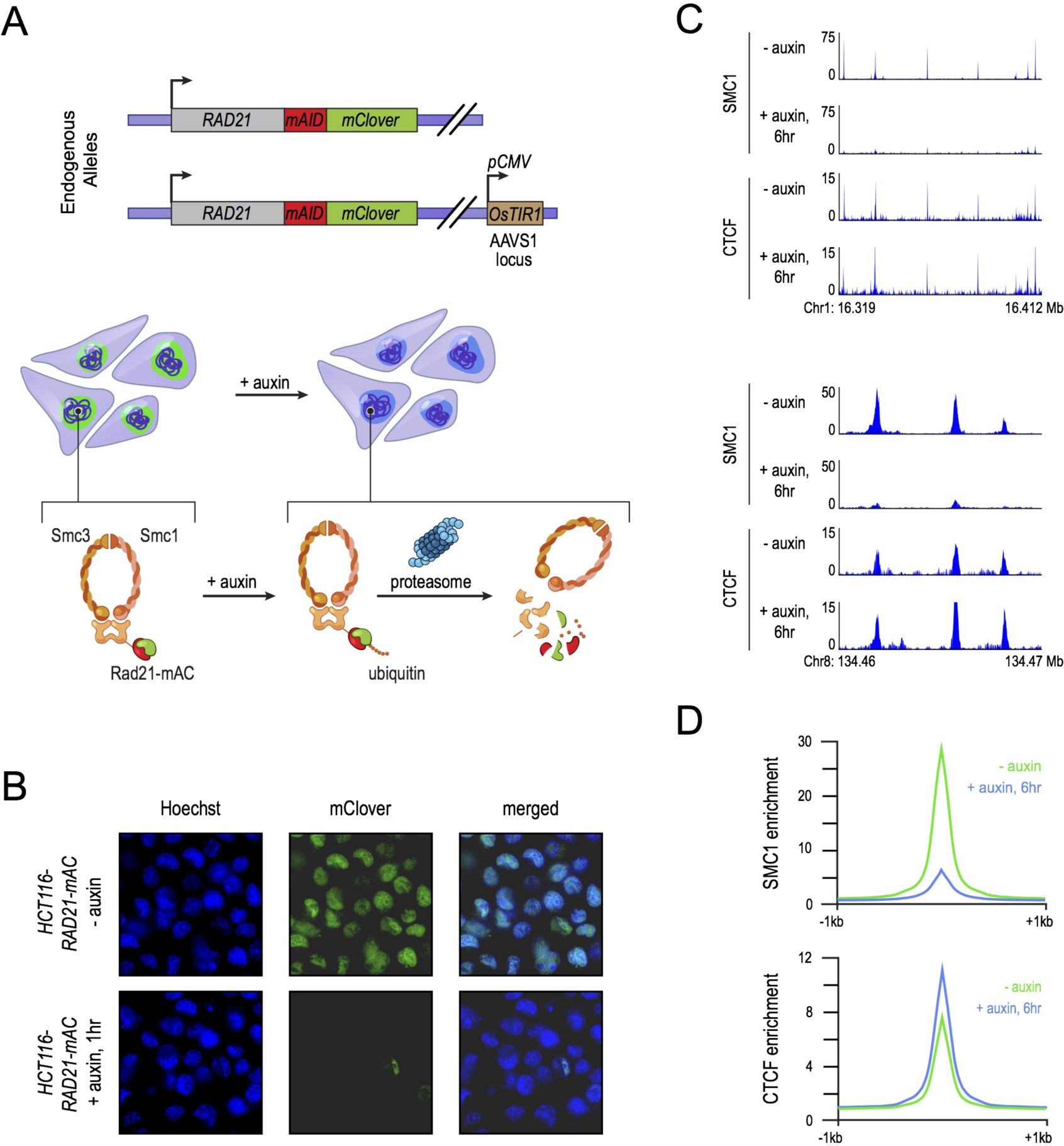
Tagging of endogenous *RAD21* with an auxin-inducible degron allows for rapid and near complete cohesin loss. (A) In HCT-116-RAD21-mAC cells (Natsume et al.), both *RAD21* alleles are tagged with auxin-inducible degrons and an mClover reporter, and the *OsTIR1* gene is integrated at the *AAVS1* locus. Addition of auxin leads to degradation of RAD21 by the proteasome. (B) Live cell imaging of HCT-116-RAD21-mAC cells after Hoechst 33342 staining to label nuclei. Nuclear mClover fluorescence corresponding to tagged RAD21 was lost after 1 hour. (See Fig. S1.) (C) SMC1 and CTCF ChIP-Seq signal with and without auxin treatment for two example loci. (D) SMC1 and CTCF ChIP-Seq enrichments averaged across ChIP-Seq peaks called by ENCODE (2012) in HCT116 cells (Natsume et al., 2016). After RAD21 degradation, the cohesin complex no longer binds to chromatin. CTCF binding is not affected.

### Histone modification patterns are unaffected by cohesin loss

We first examined the effects of cohesin degradation on key epigenomic features associated with genome folding, by performing ChIP-Seq. Using ChIP-Seq, we examined the distribution of CTCF (associated with loop anchors) and the histone modifications H3K27me3, H3K36me3, H3K27Ac, H3K4me1, H3K4me3, H3K9me3, H4K16Ac, H4K20me3, H3K79me2, and H2.AZ (associated with compartment intervals). Cohesin loss had little effect on these features (Fig. 1C,D; see Methods).

### Loop domains are rapidly lost after degradation of cohesin

We then turned to study genome folding itself, beginning with loop domains. Loops arise when two loci on the same chromosome are tethered together. (For clarity below, the loci will be referred to as “loop anchors”, the tethered pair as a “link”, and the interval between then as a “loop”.) The loop anchors are typically a pair of DNA motifs in the convergent orientation (i.e., motifs face each other) that bind CTCF and cohesin (Rao et al., 2014). Loops frequently form a contact domain—that is, an interval in which all loci exhibit higher contact frequency with one another (than random loci at similar distance along the genome sequence); this structure is called a “loop domain” (Rao et al., 2014).

To examine loop domains, we used *in situ* Hi-C (Rao et al., 2014), which combines DNA-DNA proximity ligation and high-throughput sequencing to create heat maps showing the frequency of physical contact between all pairs of loci across the genome. Loop domains are manifest in Hi-C maps as a bright “peak” pixel (indicating the link between the two loop anchors) at the corner of a bright square (indicating the presence of a contact domain).

We generated over 5 billion *in situ* Hi-C contacts from HCT-116 cells, both before (2.6B) and after (2.5B) auxin treatment. In the untreated cells, our algorithms annotated 3,170 loops, of which 2,140 were loop domains. Strikingly, the loop domains disappeared upon cohesin loss. The result was evident by visual examination (Fig. 2A, Fig. S2D–G,I–K). Moreover, the algorithms found only 9 loop domains after auxin treatment. Upon close inspection, all were found to be false positives (with 8 being due to rearrangements in HCT116 with respect to the hg19 reference genome; see Methods). (We return below to examine loops *not* associated with contact domains.)

**Figure 2:**
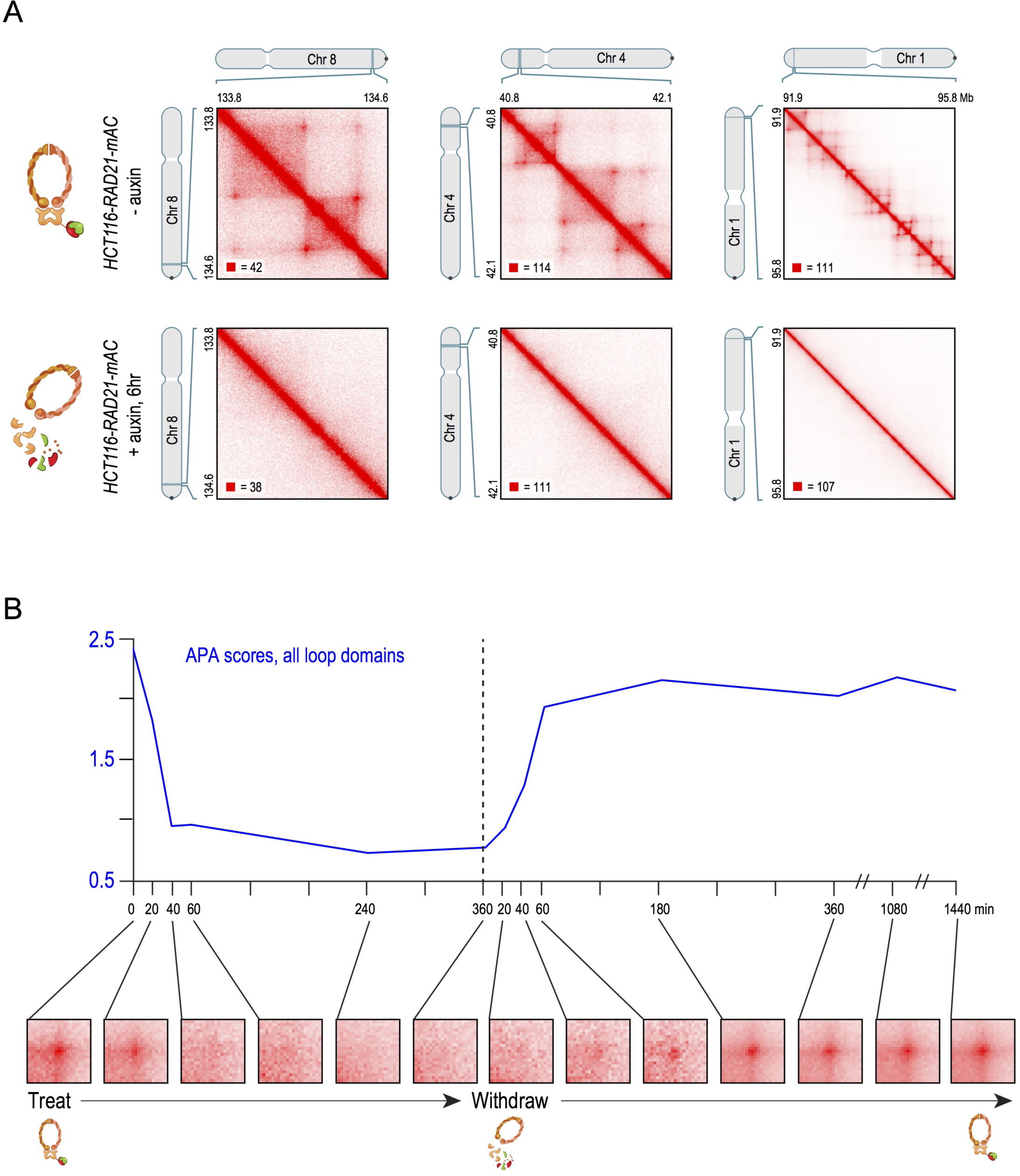
Cohesin degradation eliminates loop domains. (A) Contact matrices show that loop domains in untreated HCT-116-RAD21-mAC cells (top) disappear after auxin treatment (bottom). Three representative loci are shown (at 10kb resolution): chr8:133.8-134.6Mb (left), chr4:40.8-42.1Mb (middle) and chr1:91.9-95.8Mb (right). (B) Aggregate peak analysis (APA (Rao et al., 2014)) was used to measure the aggregate strength of the links associated with all loop domains in low-resolution Hi-C contact maps generated across a time course of auxin treatment and withdrawal. APA scores are shown on top; values greater than 1 indicate the presence of loops. APA plots for each time point are shown on the bottom; the strength of looping is indicated by the extent of focal enrichment at the center of the plot. Loop domains are rapidly lost as cohesin is degraded, and quickly restored when auxin is withdrawn (See Fig S2C).

To test whether these changes were reversible, we performed a time-course analysis in which untreated cells were exposed to auxin for six hours, after which auxin was withdrawn (Fig. 2B, Fig. S2C). Low resolution Hi-C was performed immediately before treatment, as well as at a series of time points during treatment (at 20, 40, 60, 240, and 360 minutes) and after withdrawal (at 20, 40, 60, 180, 360, 1080, and 1440 minutes). To assess whether the anchors of the loop domains seen in the pre-treatment data continued to be linked (that is, co-located in space) at each subsequent time point, we used a method called Aggregate Peak Analysis (APA) (Durand et al., 2016; Rao et al., 2014), which superimposes the signals from a set of peak pixels, thus allowing us to observe an aggregate signal even in sparse datasets where individual signals cannot be resolved (Fig. 2B). The APA signal was initially strong, but was gone by 40 minutes after treatment, and remained absent throughout the remainder of the treatment period (Fig. 2B, S2C). The disappearance of the loop-domain links closely mirrored the depletion of cohesin levels in the samples during the treatment period, as ascertained by measuring mClover fluorescence (Fig. S1). After auxin was withdrawn, the APA signal quickly increased, recovering fully by 1 hour (Fig. 2B, S2C).

These results indicate that the formation of loop domains requires cohesin; that loops and their associated contact domains rapidly disappear after the cohesin tethering the link has been degraded; and that the restoration of cohesin rescues the loops and the associated contact domains.

### Loss of cohesin is associated with stronger genome compartmentalization

Next, we examined the effects of cohesin loss on compartmentalization. Compartmentalization refers to the fact that the genome is partitioned into intervals (which can range from 6 kb to more than 5 Mb) belonging to a small number of types, such that intervals of the same type exhibit an enhanced contact frequency with one another, relative to intervals of another type (Lieberman-Aiden et al., 2009; Rao et al., 2014). Intervals are thereby assigned to two compartments (A or B, which are closely associated with open and closed chromatin, respectively) and, more finely, into six subcompartments (A1, A2, B1, B2, B3, B4). The intervals, termed “compartment intervals”, are associated with distinctive patterns of chromatin marks (Rao et al., 2014). Because loci within a compartment interval are all of the same type, they exhibit an increased contact frequency with one another and frequently form contact domains. In this case, we call the contact domain a “compartment domain.” The enhanced contact frequency between compartment intervals in the same subcompartment also gives rise to a plaid pattern in Hi-C maps (Lieberman-Aiden et al., 2009; Rao et al., 2014).

Whereas loop domains disappear entirely after cohesin loss, compartmentalization is preserved (Fig. 3A). Following auxin treatment, there is no significant change in either the compartment domains, as defined by the presence of the corresponding squares along the diagonal in the Hi-C contact map (Fig. 3B; see Methods), or in the plaid pattern, as defined by the eigenvectors of the Hi-C correlation map (Fig. 3A; mean Pearson’s r = 0.968 across all chromosomes). Our data is consistent with a previous report that genome compartmentalization is preserved after cohesin depletion (Seitan et al., 2013).

**Figure 3:**
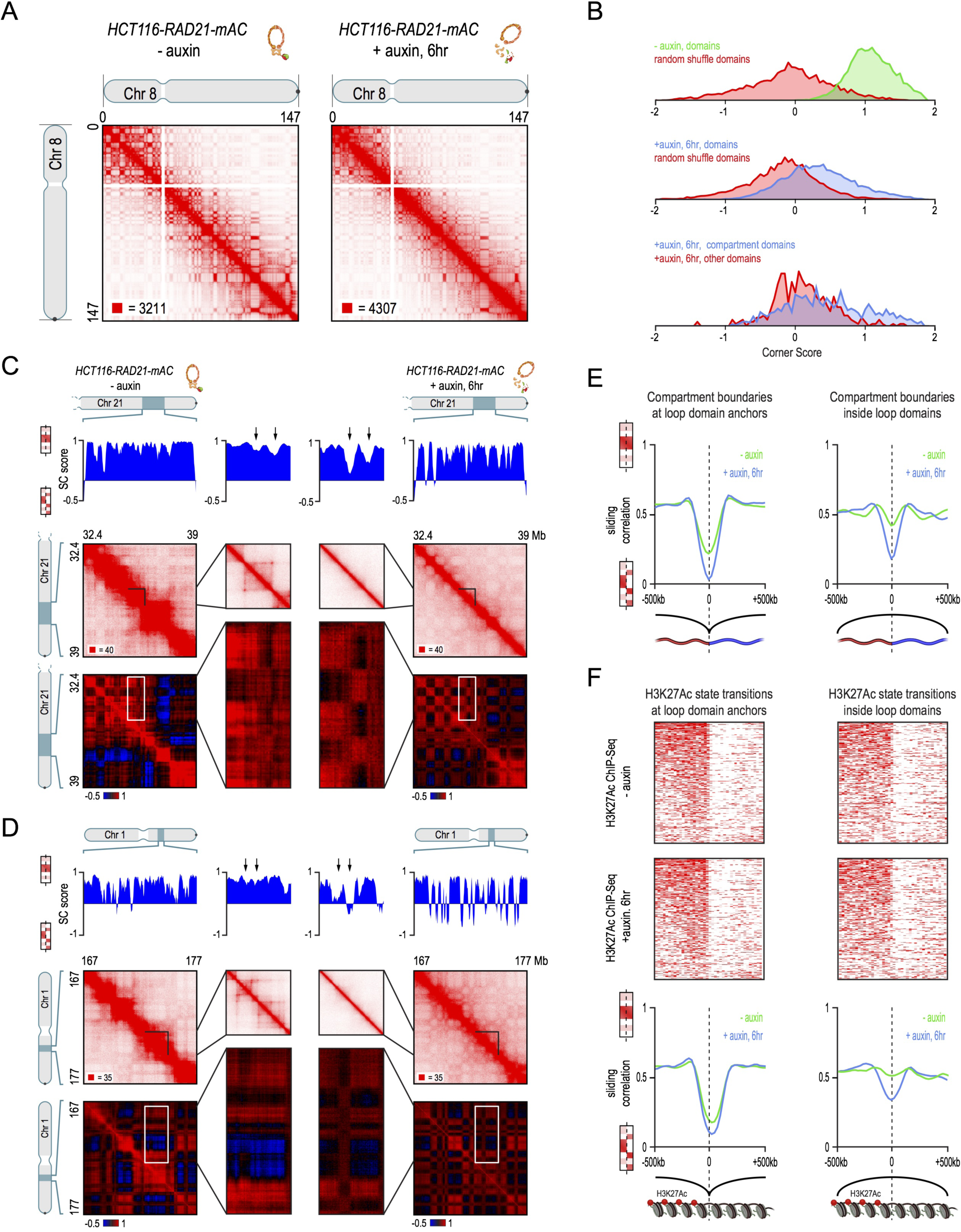
Genome compartmentalization is strengthened after cohesin degradation. (A) Contact matrices of chromosome 8 at 500kb resolution. The plaid pattern in the Hi-C map is preserved after auxin treatment, indicating that the genome compartments still form. (B) Strength of contact domains called in untreated cells versus random intervals measured using the corner score (see Methods) in untreated (top) and treated cells (middle). Contact domain strength is significantly reduced, but does not disappear. The remaining signal in treated cells comes from compartment domains (contact domains where both boundaries of the domain are also boundaries of a compartment interval), which are preserved after cohesin loss (bottom). Contact domains where both boundaries are contained completely inside a compartment interval (‘other domains’) show corner scores in treated cells comparable to random pixels. (C,D) Examples (C: chr21:32.4-39Mb and D: chr1:167-177Mb) showing that the loss of cohesin-associated loops after auxin treatment results in increased fine-scale compartmentalization. Top: Sliding correlation scores; valleys imply strong differences in long-range contact pattern observed at a locus as compared to neighboring loci, indicating a change in compartment (see Methods). Middle: Observed contact matrices. Bottom: Pearson’s correlation maps for the local region shown (see Methods). Deeper valleys in the sliding correlation score and increased plaid patterning in the observed and Pearson’s correlation maps indicate strengthened fine-scale compartment interactions after auxin treatment. Blowouts: loss of a loop domain results in strengthening of a compartment boundary spanned by the loop. Blown-out regions are indicated on zoomed out maps for both the observed (black upper triangle) and Pearson’s correlation maps (white rectangle). Observed and Pearson’s correlation maps are both shown at 25kb resolution for the zoomed out matrices and 10kb and 25kb resolution respectively for the blown-out matrices. (E) Sliding correlation scores before and after auxin treatment for compartment boundaries which either coincide with loop domain anchors (left) or are located in the interior of a loop domain (right). For compartment boundaries that lie in the interior of a loop domain in untreated cells, the difference in long-range contact pattern on opposite sides of the boundary increases greatly after cohesin treatment. (F) Sliding correlation scores before and after auxin treatment for H3K27ac boundaries in untreated cells which either coincide with loop domain anchors (left) or are located in the interior of a loop domain (right). H3K27Ac modification patterns are unchanged after auxin treatment (top and middle). For H3K27ac boundaries that lie in the interior of a loop domain in untreated cells, the difference in long-range contact pattern on opposite sides of the boundary increases greatly after cohesin treatment. This indicates that loop domains facilitate mixing of chromatin with different histone modifications.

We then examined the interaction between compartments and loop domains. Specifically, we examined the compartment boundaries (transition points between compartment intervals) that either (i) lay in the interior of a loop domain in untreated cells or (ii) coincided with a loop-domain anchor in untreated cells (Fig. 3C-E). In the former case, the correlation in the genome-wide contact pattern on opposite sides of compartment boundaries showed a much greater decrease in treated vs. untreated cells—that is, the plaid pattern across the genome became much stronger in the absence of cohesin (Fig. 3C–E). The results were similar when we examined boundaries between intervals that were enriched vs. depleted for H3K27 acetylation (which marks intervals in the “A” compartment (Rao et al., 2014)) or intervals that were enriched vs. depleted for H3K27 trimethylation (which marks intervals in the “B1” subcompartment (Rao et al., 2014)) (Fig. 3F, S4A. see Methods). These data indicate that the compartmentalization process that brings together loci with similar histone marks does not rely on cohesin. On the contrary, the strengthening of the plaid pattern after cohesin loss suggests that the formation of cohesin-dependent loop domains interferes with compartmentalization by promoting the co-localization of locus pairs with different histone modification patterns.

### Links between superenhancers are strengthened after loss of cohesin

Next, we examined loops not associated with contact domains. Whereas 1,030 such loops were annotated in untreated cells, only 72 were annotated following cohesin loss. Upon close examination, 57 were false positives (see Methods). (The loop-detection algorithms have a higher false-discovery rate after cohesin loss, since true positives are so rare.) The remaining 15 loops were much larger than those seen in untreated cells (median: 1.75 Mb, vs. 0.275 Mb). Given their large size, we found that loops could be more reliably identified in treated cells by running our peak detection algorithm (Durand et al., 2016; Rao et al., 2014) at coarser resolution (50-100 kb vs. 5-10 kb) (see Methods). This procedure identified an additional 46 loops that were confirmed by manual inspection (Fig. 4A, see Methods). After including these additional loops, the size difference between the 61 “cohesin-independent loops” and the cohesin-associated loops was even more dramatic (Fig. 4C, median size: 23.15 Mb).

**Figure 4:**
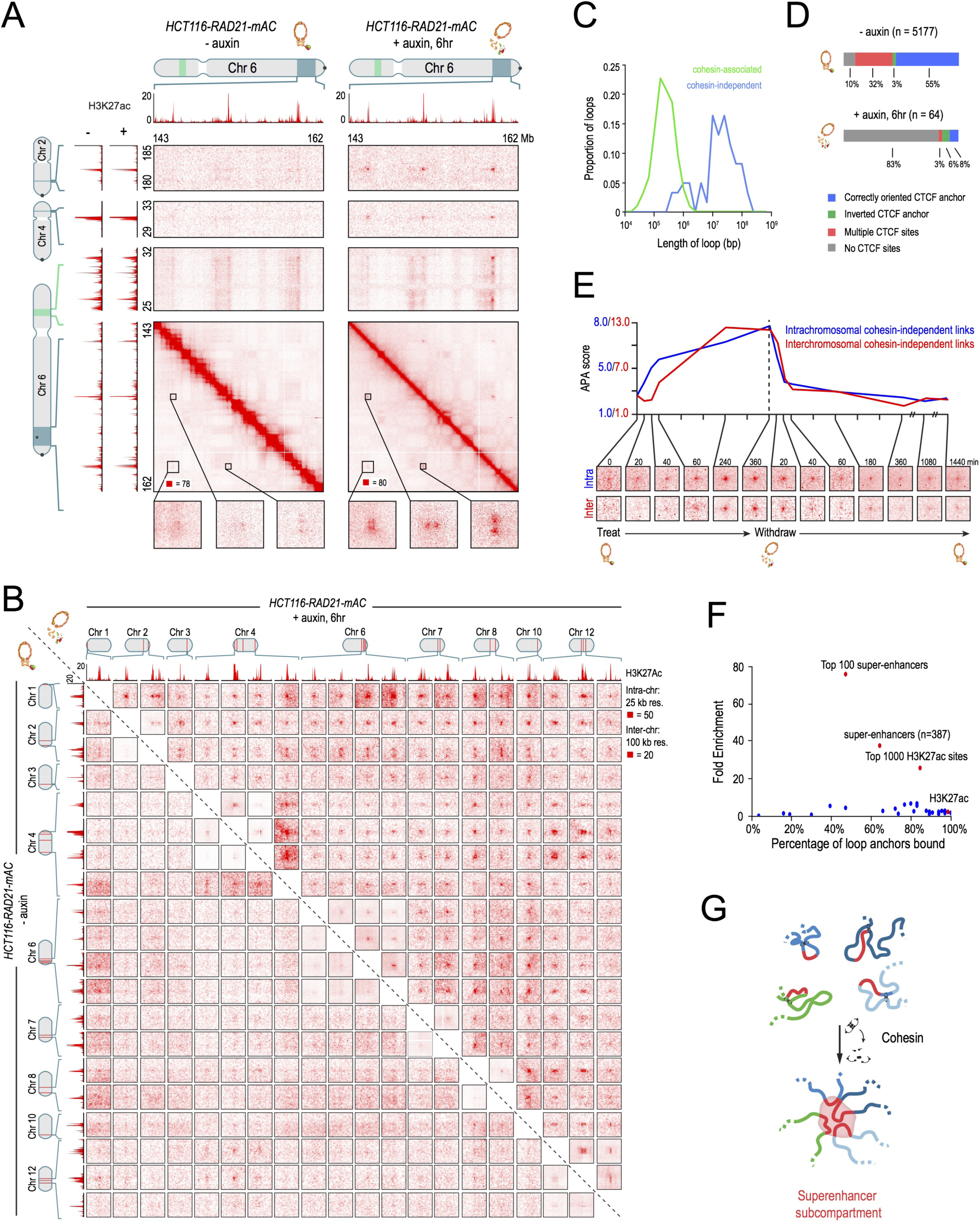
Cohesin loss causes superenhancers to co-localize, forming hundreds of links within and across chromosomes. (A) Examples of a network of intra- and interchromosomal cohesin-independent links between superenhancers on chr6, chr4, and chr2. H3K27 acetylation does not change with auxin treatment, but cohesin-independent links are significantly strengthened upon treatment. Intrachromosomal matrices are shown at 25kb (on-diagonal) and 50kb (off-diagonal) resolutions; interchromosomal matrices are shown at 100kb resolution. Maximum color intensities are 28 reads for the offdiagonal intrachromosomal matrices and 20 reads for the interchromosomal matrices. (B) The interactions between 20 cohesin-independent loop anchors spread across 9 chromosomes are shown before (lower triangle) and after (upper triangle) auxin treatment. Each matrix shows a 2 Mb by 2 Mb matrix centered on the respective anchors. Intrachromosomal interactions are shown at 25kb resolution with a maximum intensity of 50 reads; interchromosomal interactions are shown at 100kb resolution with a maximum intensity of 20 reads. The anchors are strongly enriched for H3K27 acetylation both before and after auxin treatment. (The ChIP-Seq track is shown at 25kb resolution.) Cohesin loss causes the anchors to form a clique, with focal interactions seen between nearly all pairs of loop anchors, regardless of whether they lie on the same chromosome. (C) Length distribution of cohesin-mediated loops (green) versus cohesin-independent loops (blue). (D) CTCF binding patterns at cohesin-associated (top) versus cohesin-independent loop anchors (bottom). Only cohesin-associated loops obey the convergent rule. (E) APA for intrachromosomal (blue) and interchromosomal (red) cohesin-independent links across a time course of auxin treatment and withdrawal. APA scores are shown on top and APA plots for each time point are shown on the bottom. Cohesin-independent links are rapidly strengthened as cohesin is degraded and weaken as cohesin is restored. (F) Percent of cohesin-independent loop anchors bound versus fold enrichment for 36 DNA-binding proteins and histone modifications. Superenhancers are strongly enriched at cohesin-independent loop anchors. (G) Under normal circumstances, short-range contacts between superenhancers and neighboring loci are facilitated by loop domain formation by cohesin. Upon cohesin loss, superenhancers cluster even when located on different chromosomes to form a subcompartment.

We then sought to understand the basis of these “cohesin-independent loops.” We found that cohesin-independent loops do not demarcate the boundaries of contact domains (0 of 61, or 0%; vs. 2140 of 3170, or 68%, for cohesin-associated loops). Remarkably, we observed many cohesin-independent loop anchors form links with each another – manifest as focal peaks in the Hi-C heatmap – even when the anchors reside on different chromosomes (Fig. 4A, S5F,G). In total, we identified 203 such interchromosomal links. Notably, we observed large cliques forming between the anchors of the cohesin-independent loops (Fig. 4B). These features are not seen for cohesin-associated loops (Fig. S2H).

The anchors of cohesin-independent loops also exhibit very different patterns of protein binding. The proportion that binds CTCF is much lower (20% vs. 90% for cohesin-associated loops; Fig. 4D, S5A). Moreover, there is no tendency for the CTCF motifs to point into the loop (5 of 9 (56%) point inward vs. 2770 of 2919 (95%) for cohesin-associated loops.

Notably, the cohesin-independent loop anchors are highly enriched for superenhancers (also called stretch enhancers), which are regions of the mammalian genome containing a very high density of enhancer elements, and which are marked by extremely high levels of H3K27 acetylation (Hnisz et al., 2013; et al., 2013; Whyte et al., 2013). We found that 41 of the 64 cohesin-independent loop anchors overlapped with the 387 superenhancers in HCT116 cells—a 37.5-fold enrichment, which was highly statistically significant (p<10^−15^). For the 100 strongest superenhancers, the enrichment was even more pronounced (76-fold, 30 of 64 anchors, p<10^−15^; Fig. 4F, S5C). Interestingly, loops and links between superenhancers could be seen in the untreated cells as well, but were much weaker (Fig. 4A,B,E, S5B,D–G, see Methods).

In many respects, the cohesin-independent loops resemble the superloops we previously observed on the inactive X chromosome (Darrow et al., 2016; Rao et al., 2014). These superloops are very large (up to 77Mb), the intervals they span do not form contact domains, and their anchors are marked by H3K27 acetylation (Fig. S6A–C).

Our results highlight two distinct mechanisms that guide genome folding. The first mechanism is the cohesin-dependent formation of loop domains. We (Sanborn et al., 2015) and others (Alipour and Marko, 2012; Fudenberg et al., 2016; Nasmyth, 2001) recently hypothesized that the underlying physical process is the formation of loops by extrusion. The data presented above are consistent with several models, including the possibility that loop domains form when a cohesin-based extrusion complex, which comprises two physically tethered subunits, binds chromatin at a particular location; subsequently, the subunits slide in opposite directions until they arrive at a bound CTCF protein. Thus, the disappearance of cohesin eliminates all loop domains without influencing CTCF binding.

The second mechanism is the segregation of genomic intervals into a small number of distinct spatial compartments. Our data indicate that this process does not depend on cohesin. Instead, regions with similar histone modifications may tend to co-localize with one another, perhaps through a process similar to phase separation of a block copolymer (Hnisz et al., 2017; Di Pierro et al., 2016). Our data suggest that cohesin-mediated loop domain formation partially interferes with the process of compartmentalization, for example by perturbing the phase separation. In the absence of cohesin, compartmentalization becomes sharper.

The two mechanisms also suggest an explanation for the cohesin-independent loops—namely, that the links between these anchors represent compartmental co-segregation (rather than cohesin mediated tethers) between small intervals containing H3K27-acetylated superenhancers (Fig. 4G). This explanation accounts for why these links can exist both within and between chromosomes, why the links are weaker in the presence of cohesin, and why the anchors form large cliques. It is possible that short genomic intervals decorated by other chromatin marks may, in certain cases, co-segregate in a similar fashion.

### Molecular dynamics simulations integrating extrusion and compartmentalization can recapitulate Hi-C experimental results

To test the hypothesis that the Hi-C contact maps we observed are consistent with the presence of two distinct folding mechanisms, we modeled a 2.1 Mb region on chromosome 3 as a block copolymer consisting of two types of chromatin, A or B, determined by the contact pattern observed in the treated Hi-C map; and containing CTCF binding sites whose position and strength were derived from CTCF and SMC1 ChIP-Seq tracks, and whose orientation was determined by examining the human genome reference. We used molecular dynamics simulations to examine the behavior of this polymer in a solvent containing extrusion complexes (thus modeling loop extrusion (Alipour and Marko, 2012; Fudenberg et al., 2016; Sanborn et al., 2015)), and in the presence of attractive forces between like monomers (thus modeling compartmentalization (Barbieri et al., 2012; Di Pierro et al., 2016)). The resulting ensemble was used to create an *in silico* contact map for the region. (These simulations build on earlier work that predicted contact maps resulting from loop extrusion using CTCF ChIP-Seq tracks and the reference genome as the only inputs (Sanborn et al., 2015).)

We found that the resulting contact maps accurately recapitulated the plaid pattern of compartment interactions seen both in the presence and in the absence of cohesin. They also recapitulated the positions of loop domains in the untreated cells, and their disappearance after cohesin degradation (Figure 5A,B). The simulations also illustrate the change in long-range contact pattern that is seen when a loop spans a compartment boundary.

**Figure 5:**
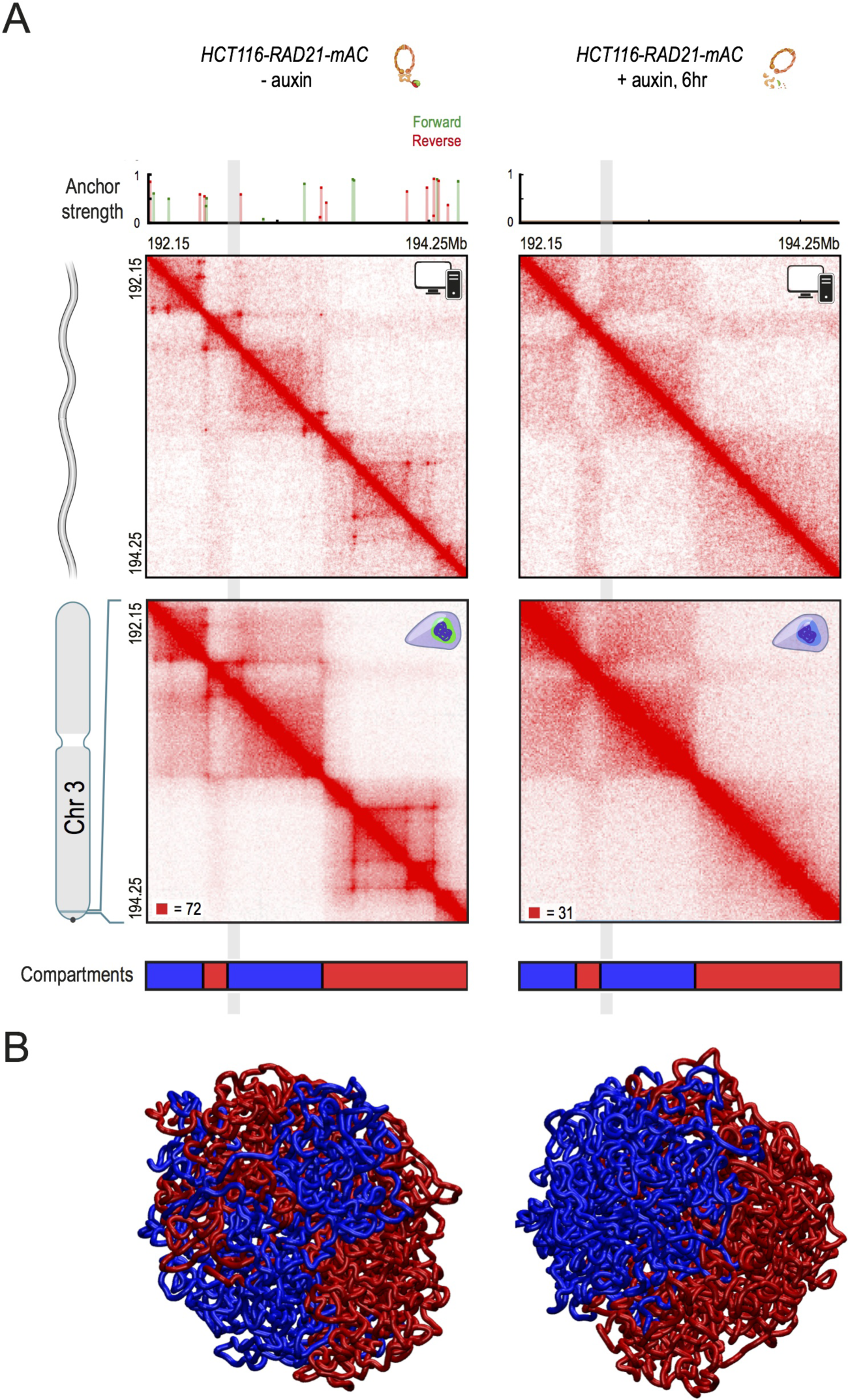
Molecular dynamics simulations combining extrusion and compartmentalization can recapitulate Hi-C experimental results. (A) We use loop extrusion and compartmentalization to simulate a 2.1 Mb region on chromosome 3 in HCT-116 RAD21-mAC cells before (left) and after (right) auxin treatment. SMC1 ChIP-Seq signals are normalized and converted into binding probabilities for the simulated extrusion complex. Each peak is assigned a forward (green) or reverse (red) orientation based on the corresponding CTCF motif. Hi-C contact patterns in the treated map were used to determine the positions of compartment intervals (red and blue). The simulations yield an ensemble of polymer configurations. We show contact maps from the simulated ensemble (top) and from the corresponding Hi-C experiments (bottom). The simulations accurately capture the positions of loops and domains, as well as the loss of loop domains after the depletion of cohesin. In addition, our simulation accurately captures compartmentalization patterns seen before and after auxin treatment. Notably, one of the loop domains spans multiple compartment intervals; the loci between the boundary of one of the compartment intervals and the loop anchor are highlighted (grey). (B) Examples of globules from simulations of compartmentalization with extrusion (left) and without (right). Notably, the globule without extrusion shows stronger segregation of compartment types.

### Cohesin loss results in strong down-regulation of genes near superenhancers, but does not bring about widespread ectopic activation

Finally, we sought to investigate the role of cohesin in the regulation of gene expression. Cohesin has been proposed to facilitate interactions between enhancers and promoters (Chien et al., 2011; Kagey et al., 2010; Merkenschlager and Odom, 2013; Phillips-Cremins et al., 2013). Loop domains are thought to regulate this process by preventing enhancers from forming ectopic interactions with targets that lie in a different loop domain (Dixon et al., 2016; Dowen et al., 2014; Flavahan et al., 2016; Krijger and de Laat, 2016; Lupiáñez et al., 2015; Narendra et al., 2015). We therefore characterized the effects of cohesin loss on nascent transcription by performing precision nuclear run-on sequencing (PRO-Seq) in treated and untreated HCT116 cells (Engreitz et al., 2016; Kwak et al., 2013) (Fig. 6A). We chose an early timepoint - 6 hours after auxin treatment – with the aim of examining direct consequences, rather than indirect effects due to changes in cell state.

**Figure 6:**
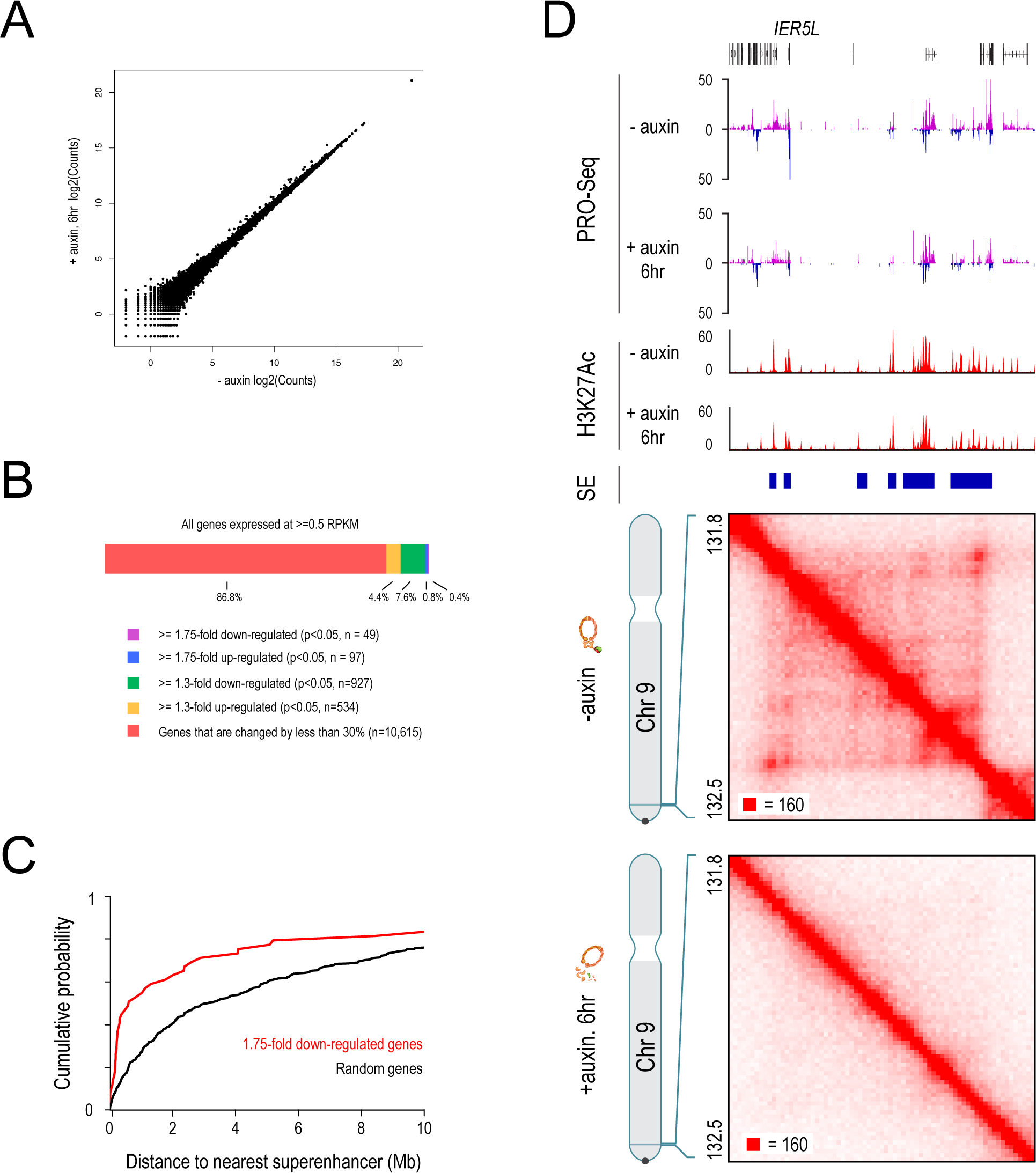
Cohesin degradation results in strong down-regulation of genes near superenhancers but does not result in widespread ectopic gene activation. (A) Scatter plot of gene-wide PRO-Seq counts in RAD21-mAC cells before (x-axis) and after (y-axis) treatment. (B) Genes that are expressed in untreated cells rarely undergo substantial changes in expression level after cohesin loss. (C) Cumulative probability distributions of distances to the nearest superenhancer for 1.75-fold down-regulated genes after auxin treatment (red) versus random genes (black). (D) An example of a strongly down-regulated gene near a superenhancer. In untreated cells, a series of cohesin-associated loops form between the *IER5L* promoter and nearby superenhancers. Upon auxin treatment, these loops are lost and *IER5L* expression is 2.6-fold down-regulated. (Note that, unlike this example, many of the genes that are strongly down-regulated after treatment do not exhibit visible promoter-enhancer looping in untreated cells.)

**Figure 7:**
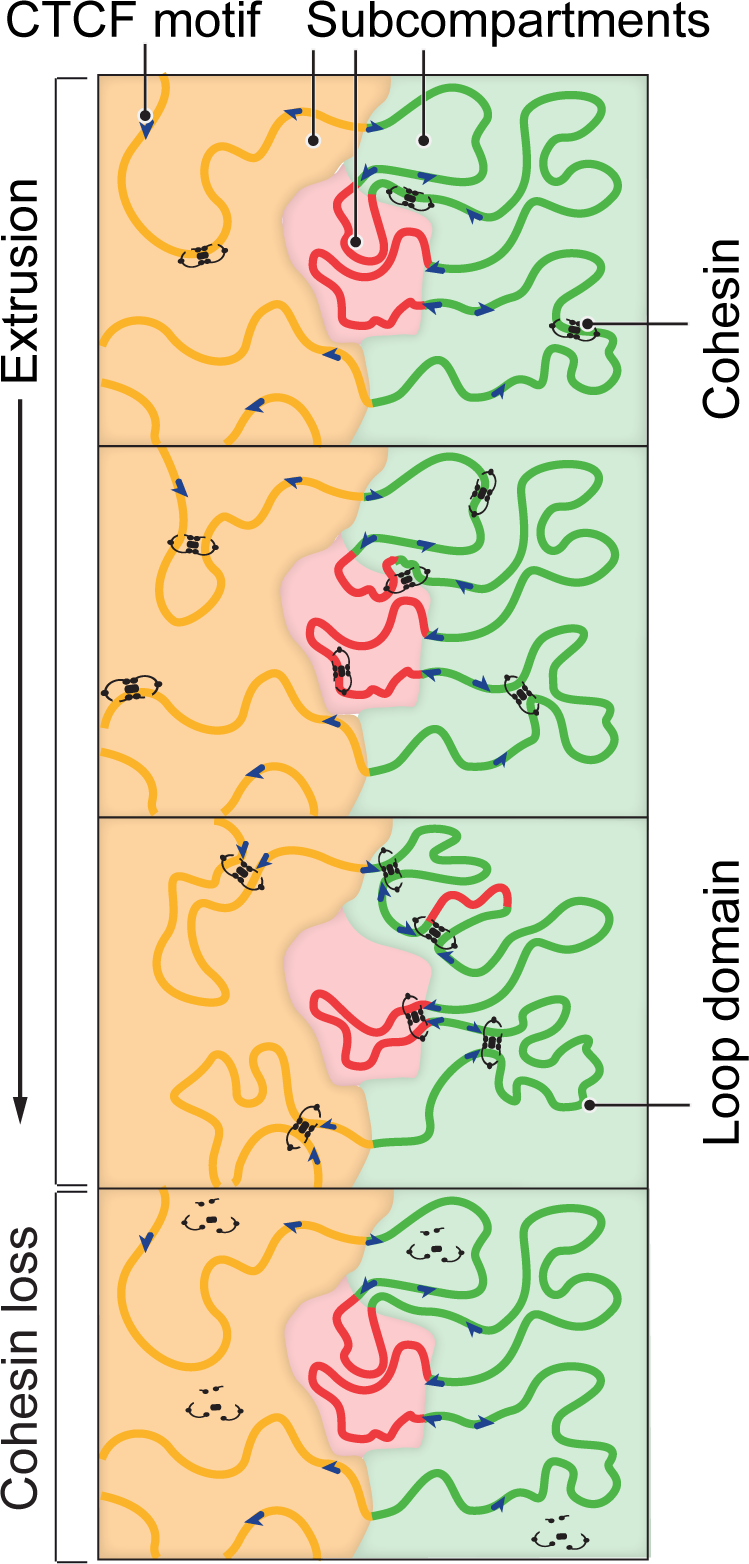
Extrusion and compartmentalization interact to shape the spatial organization of the genome inside the nucleus. Intervals of chromatin with similar patterns of histone modification colocalize in nuclear subcompartments. Loop extrusion facilitates short-range contacts between nearby loci as the two subunits of the cohesin complex translocate in opposite directions on chromatin until they halt at CTCF motifs in the convergent orientation, thus forming a loop domain. Loop domains represent dynamic structures that are maintained by cohesin; only a subset of them may be present at any given time. When these anchor motifs span multiple compartment intervals, loop extrusion interferes with compartmentalization by facilitating contacts between loci in different compartments. Loss of cohesin leads to the disappearance of loop domains and to a closer correspondence between genome compartmentalization patterns and histone modification patterns.

To look for signs of ectopic activation, we examined the 14,853 genes that were not expressed (RPKM<0.5) in untreated cells. Of these genes, 1% (216) were ectopically activated after treatment (p<0.05, >30% change in RPKM, RPKM>0.5 in treated cells). In addition, 7% of these genes (1063) exhibited “leaky” transcription in treated cells: a larger PRO-Seq signal (p<0.05, >1.3 fold change difference) that fell short of the threshold for an expressed gene (i.e., RPKM was still below 0.5). These findings imply that, while cohesin plays a role in preventing ectopic activation, most genes remain inactivated even in the absence of cohesin and loop domains.

We next looked for changes in the 12,222 genes that were expressed (RPKM>0.5) in untreated cells (Fig. 5B). Here again, most genes (87%, 10,615) exhibited similar levels of transcription after cohesin degradation (RPKM changed by less than 30%). The remaining genes (13%, 1607) showed a larger transcriptional effect (p<0.05, >30% change in RPKM). Stronger effects were seen, but less frequently: 64 genes (0.5%) showed a 2-fold change, and 2 genes showed a 5-fold change (Fig. 6B). While the quantitative impact may seem modest, we note that such changes can have important biological impacts—for example, two-fold increases in expression of receptor tyrosine kinase genes can double cell growth rates and play important roles in cancer.

Of genes that exhibited a strong change in transcription, more were downregulated than upregulated (61% vs. 39%)—suggesting that cohesin-associated loops may both facilitate activation of promoters by their distal enhancer elements and block activation by inappropriate enhancers, with the former being somewhat more common.

We wondered how cohesin facilitated these promoter-enhancer contacts. We noticed that many of the genes that were downregulated (by >1.75-fold) were located within 500kb of superenhancers (23 of 49, 4.8-fold enrichment, Fig. 6C,D; S8A–B). Of these genes, 29% were located with 500kb of one of the top 100 superenhancers (8.5-fold enrichment). Strikingly, these superenhancers were often located at the anchors of the cohesin-independent links seen in treated cells (8 of 19, a 13.7-fold enrichment).

Taken together, our results indicate that, in the absence of cohesin, superenhancers associated with the downregulated genes exhibit a strong tendency to form links with one another. By contrast, in the presence of cohesin, the majority of these superenhancers were located in the interior of cohesin-associated loop (in 13 of 19 cases) and the long-distance cohesin-independent links were much weaker. This is consistent with our earlier finding that compartmental co-segregation appears to have less influence on long-range contact patterns when compartment boundaries lie in the interior of a cohesin-associated loop.

The above results are interesting from the standpoint of transcriptional regulation. Our data suggest a model where cohesin-associated looping, by increasing the frequency of contact between loci within loop domains and by disturbing patterns of compartmentalization, facilitates mixing between elements (such as genes and superenhancers) that would otherwise be segregated. In this model, compartmentalization and extrusion – through independent mechanisms – interact to shape transcription.

## DISCUSSION

Here, we use extremely high resolution maps of a human colon cancer cell line before and after cohesin degradation to demonstrate that the cohesin protein RAD21 is required for the establishment and maintenance of loop domains genome-wide. After cohesin loss, we also find that: (i) histone marks are unchanged; (ii) compartment structure is strengthened in the absence of cohesin, as loop domains spanning multiple compartment intervals lead to mixing among loci in different compartments; (iii) only a small subset of genes exhibit large changes in transcription level. Finally, we identify a class of cohesin-independent loops and links connecting superenhancer loci on different chromosomes.

### Comparison with other studies

It is important to set our results in the context of other studies related to loop formation. While there is clear agreement that deletion of individual CTCF binding sites can result in a loss of cohesin binding and can abolish loops and contact domains (Guo et al., 2015; Narendra et al., 2015; Sanborn et al., 2015; de Wit et al., 2015), there have been conflicting reports about the effects of depleting cohesin or CTCF.

In contrast to the high-resolution Hi-C studies reported here, early Hi-C studies of cohesin depletion, using both gene knockouts and proteolytic cleavage, reported that contact domains remained (Seitan et al., 2013; Sofueva et al., 2013; Zuin et al., 2014). The discrepancy may be due to (i) the fact that low resolution Hi-C analysis cannot not distinguish between loop domains, which are sensitive to cohesin depletion, and compartment domains, which are not; and (ii) the possibility that the cohesin depletion was incomplete. A recent preprint sought to extract more information from these early studies by using genome-wide aggregate signals (rather than, as here, signals from individual loops) (Schwarzer *et al.* 2016). It found that the data continue to show the presence of loops—a conclusion that we confirmed through independent analysis (Figure S3). These results suggest that the earlier experiments were performed in the setting of incomplete cohesin loss (in the case of cleavage) or incomplete cell purification (in the case of gene knockouts), a possibility that the early studies acknowledge.

Two groups have also recently posted preprints exploring the effects of CTCF depletion using higher-resolution Hi-C maps. One of these studies reported that CTCF depletion leads to the disappearance of contact domains (Nora et al., 2017), while the other study reached the opposite conclusion (Kubo et al., 2017). Our data for cohesin depletion is consistent with the former preprint, because CTCF depletion would be expected to lead to cohesin sliding no longer being arrested at CTCF sites. The reason for the discrepancy between these two preprints is not immediately clear, although it might be due to lack of complete CTCF degradation in one of the studies.

Two recent studies have also sought to examine the effects of depletion of *NIPBL,* which encodes a cohesin loader protein. They too report opposite conclusions. The (Schwarzer et al., 2016) preprint reported, on the basis of new experiments, that the genome-wide aggregate signal from loops is absent following *NIPBL* deletion. By contrast, a recent publication reported the continued presence of individual loop domains after the near-complete depletion of *NIPBL,* although these loop domains were abnormally small (Haarhuis et al., 2017). This discrepancy could be explained by the fact that the former paper analyzed the genome-wide aggregate signal, which (i) can fail to observe looping if the distribution of loop sizes shifts dramatically away from expectations and (ii) is not sensitive to the cohesin-independent loops between superenhancers that we report, which are much larger than previously reported autosomal loops. In any case, it is unclear what effect NIBPL depletion would be expected to have on loop domain formation because, although *NIPBL* facilitates cohesin loading, it may not be essential for cohesin loading. Indeed, cohesin loading independent of NIPBL has been observed *in vitro*, albeit at low efficiency (Davidson et al., 2016; Murayama and Uhlmann, 2014; Stigler et al., 2016).

Finally, an exciting recent experiment demonstrated that deletion of WAPL, a cohesin antagonist that is responsible for the disassociation of cohesin from chromatin, results in the formation of thousands of new loops and loop domains, which are larger than those found when WAPL is intact (Haarhuis et al., 2017). Our results are consistent with these findings.

### The speed of loop extrusion

Our work demonstrates that loop domains disappear shortly after auxin-induced cohesin loss, and that they are restored within 40 minutes of auxin withdrawal. These findings imply that cohesin is required both for the formation of loop domains, and for their maintenance. Notably, the observation that loop domains disappear within minutes of cohesin loss argues against a model in which they represent stable states of chromosome condensation.

In the loop extrusion model, two physically tethered subunits bind chromatin at a single site and then slide in opposite directions along the 1D contour of the chromosome. In the context of this model, the rapid formation of loops between anchor loci that are megabases apart implies that each subunit of this extrusion complex slides hundreds of base pairs per second. This estimate is a lower bound, as it ignores the time needed for auxin to disappear from individual cells, the time needed for transcription of RAD21 protein, and the time needed for the tripartite cohesin ring complex to re-form and be loaded on chromatin. This estimated translocation rate is similar to estimates obtained studying the SMC complex in *B. subtilis* as it travels from the origin of replication to the terminus [500-1000 bp/s] (Wang et al., 2017).

Notably, the estimated rate bears on the protein motors involved when extrusion complex subunits slide. For instance, it is possible that cohesin itself serves as a motor during the extrusion process. However, single molecule studies of cohesin translocation *in vitro*, have yielded an estimated sliding rate of only 1-2bp per second on chromatin (Davidson et al., 2016; Kanke et al., 2016; Kim and Loparo, 2016; Stigler et al., 2016). This suggests that cohesin alone is unlikely to be the principal source of translocase activity.

Similarly, it was recently shown, *in vitro*, that RNA polymerase II can push cohesin rings along DNA (Davidson et al., 2016). It has also been shown that cohesin accumulates at sites of convergent transcription in yeast and in mammals (in the absence of CTCF and WAPL), raising the possibility that polymerase might also play a role in cohesin translocation *in vivo* (Busslinger et al., 2017; Lengronne et al., 2004). However, current estimates of the elongation rate of RNA polymerase (9-90 bp/s) suggest that polymerase-driven sliding would be much slower than the above estimate (Jonkers and Lis, 2015). This suggests that other translocases must be involved in loop extrusion.

Going forward, the loop extrusion model will be greatly clarified by detailed studies of cohesin translocation, and of the motor proteins involved.

### The interplay between loop extrusion and compartmentalization

Our results are consistent with the presence of two distinct mechanisms of genome folding: (i) extrusion, which results in the formation of loop domains, and which is entirely dependent on cohesin; and (ii) compartmentalization, which results in the formation of compartment domains and in the cosegregation of similar compartment intervals both within and across chromosomes. The latter mechanism remains after cohesin depletion (Seitan et al., 2013)

Using our high-resolution contact maps, we are also able to examine the ways in which these two mechanisms interact. It is commonly thought that compartment intervals are typically megabases in length, and are subdivided into smaller domains in a hierarchical fashion (Dixon et al., 2016; Fraser et al., 2015; Gibcus and Dekker, 2013; Gorkin et al., 2014; Nora et al., 2013; Sexton and Cavalli, 2015; Shachar and Misteli, 2017). Here, we demonstrate that compartment intervals can be as short as tens of kilobases, and can overlap loop domains in complex ways. For instance, we observe numerous examples of loop domains spanning multiple compartment intervals.

Moreover, we find that cohesin-mediated loop extrusion, by facilitating contacts between all loci in the loop, can enhance the contact frequency between loci that would ordinarily lie in different subcompartments. Thus, when multiple compartment intervals lie in a single loop, the long-range contact pattern seen for each locus is a mixture of the pattern that would ordinarily be seen for loci in the corresponding subcompartment, and the pattern seen for other loci in the loop. This mixing disappears upon cohesin depletion (Fig. 7). Similarly, deletion of WAPL appears to increase the processivity of the loop extrusion complex, and thereby to increase the size of loops (Busslinger et al., 2017; Haarhuis et al., 2017; Tedeschi et al., 2013). Consistent with our observations, these larger loops are associated with extensive mixing, which largely obscures the long-range compartment patterns (Haarhuis et al., 2017).

A dramatic example of this effect is seen in the case of superenhancers, which are short intervals of chromatin showing extremely high H3K27 acetylation. In the presence of cohesin, superenhancer loci typically lie within cohesin-dependent loops, which limits their ability to co-segregate by means of genome compartmentalization. Following cohesin loss, superenhancer loci are more readily able to co-localize via compartmentalization, forming loops and links in large cliques both within and across chromosomes.

Notably, the loops and links between pairs of superenhancers increase in strength rapidly following cohesin loss, reaching a plateau within hours. The speed at which these small genomic elements are able to co-localize, even when located on different chromosomes, implies that compartmentalization is capable of inducing focal interactions genome-wide at rates comparable to those of loop-domain formation.

What physical mechanisms might be involved in cohesin-independent folding in the nucleus? Studies spanning multiple organisms have observed strong correlations between histone modification patterns and long-range contact patterns in Hi-C maps (Dixon et al., 2012; Lieberman-Aiden et al., 2009; Rao et al., 2014; Ryba et al., 2010; Sexton et al., 2012; Zhu et al., 2013). However, it is currently unclear whether histone modifications drive the formation of compartments; whether compartmentalization drives the deposition of histone modifications; whether the two processes reinforce each other; or whether the two processes are driven by a third mechanism. The fact that our experiments find that cohesin loss does not affect histone modifications, but does cause long-range contact patterns to better match patterns of histone marks, is more consistent with histone patterns governing genome compartmentalization, rather than the reverse. Mechanistically, this process could be facilitated by phase separation of nucleosomes with similar marks (Hnisz et al., 2017; Di Pierro et al., 2016) or binding of reader proteins that target modified histones to specific locations in the nucleus (Barbieri et al., 2012; Isono et al., 2013; van de Werken et al., 2017; Wijchers et al., 2016). Of course, it remains possible that independent mechanisms may shape both processes.

### The interplay between cohesin and gene regulation

Many studies have proposed that cohesin facilitates interactions between enhancers and promoters, thereby upregulating the transcription of many genes (Kagey et al., 2010; Merkenschlager and Odom, 2013; Phillips-Cremins et al., 2013). Moreover, studies have also suggested that loop domains formed between CTCF and cohesin binding sites create insulated regulatory neighborhoods—partially protecting genes with a loop domain from the influence of enhancers outside the domains (Dowen et al., 2014; Flavahan et al., 2016; Lupiáñez et al., 2015; Narendra et al., 2015). Our study, combining rapid depletion of cohesin and measurement of nascent transcription using PRO-Seq, allows us to more clearly dissect the direct effects of cohesin loss on transcription.

We find that a very small set of genes becomes strongly downregulated after cohesin loss. These genes tend to be close to superenhancers, suggesting that cohesin indeed serves to facilitate interactions between enhancers and promoters.

However, most genes are not strongly increased or decreased by cohesin loss. This suggests that cohesin-dependent loop domains themselves play at most a modest role in facilitating or disrupting interactions between promoters and enhancers. Of course, we cannot dismiss modest effects on overall level of transcription as unimportant. For example, modest increases in the expression of receptor tyrosine kinase genes can have meaningful effects on cell proliferation. Moreover, it is possible that removal of loop domains through other mechanisms (for example, depletion of CTCF) could have a different effect on gene activation, since, for example, it would not entirely eliminate the extrusion mechanism.

In conclusion, we find that the efficient degradation of cohesin provides an illuminating window on the dynamics of genome folding as a whole, eliminating certain classes of loops and domains while leaving others intact. Systematic manipulation of the underlying machinery should help clarify the basis of genome architecture going forward.

## Author Contributions

S.S.P.R. and E.L.A conceived this project and designed all experiments. S-C.H. and B-T.S.H., and A.D.O. performed Hi-C experiments. J.E. aided with the design of the PRO-Seq experiments and J.E. and E.M.P. performed PRO-Seq experiments. K-R.K-K., S.E.J., and S-C.H. performed ChIP-Seq experiments. A.L.S. performed simulations. I.D.B. performed microscopy experiments. S.S.P.R., J.E., X.H., M.S.S., B.E.B., R.C., E.S.L, and E.L.A. analyzed data. S.S.P.R., E.S.L., and E.L.A. prepared the manuscript with input from all authors.

## Acknowledgments

This work was supported by a Paul and Daisy Soros Fellowship, a Fannie and John Hertz Foundation Fellowship and a Stanford Medical Scholars Fellowship to S.S.P.R., an NIH New Innovator Award (1DP2OD008540-01), an NSF Physics Frontier Center Grant (PHY-1427654, Center for Theoretical Biological Physics), the NHGRI Center for Excellence for Genomic Sciences (HG006193), the Welch Foundation (Q-1866), an NVIDIA Research Center Award, an IBM University Challenge Award, a Google Research Award, a Cancer Prevention Research Institute of Texas Scholar Award (R1304), a McNair Medical Institute Scholar Award, an NIH 4D Nucleome Grant (U01HL130010), NIH Encyclopedia of DNA Elements Mapping Center Award (UM1HG009375), and the President’s Early Career Award in Science and Engineering to E.L.A. We are grateful to Masato Kanemaki for sharing the HCT-116 RAD21-mAID-mClover cell line. We thank Roger Kornberg, Peter Geiduschek and Miriam Huntley for their thoughtful comments on this manuscript. The experiments in this study were informed by discussions with E. Nora regarding his experiments on domain structure after CTCF degradation, and we thank him and his collaborators.

## CONTACT FOR REAGENT AND RESOURCE SHARING

All requests for information, reagents and resources should be directed to the Lead Contact, Erez Lieberman Aiden.

## EXPERIMENTAL MODEL AND SUBJECT DETAILS

### HCT-116 cells

We obtained HCT-116-CMV-OsTir1 and HCT-116-RAD21-mAID-mClover cells (HCT-116 RAD21-mAC) from (Natsume et al., 2016). The cells were cultured in McCoy’s 5A medium supplemented with 10% FBS, 2 mM L-glutamine, 100 U/ml penicillin, and 100ug/ml streptomycin at 37C with 5% CO2. Degradation of the AID-tagged RAD21 was induced by the addition of 500uM indole-3-acetic acid (IAA; Sigma Aldrich). For our standard *in situ* Hi-C, ChIP-Seq, and PRO-Seq experiments on untreated cells and cells treated for 6 hours, medium was aspirated at t=0, and either replaced with fresh medium (untreated) or medium containing 500uM IAA. The cells were then washed, trypsinized and processed for downstream experiments at t=6hrs.

For our time course experiments, cells were treated with 500uM IAA and crosslinked with 1% formaldehyde directly in wells of a 6-well plate at various time points after treatment (20, 40, 60, 240, 360 minutes). For the auxin withdrawal experiments, after 6 hours treatment with 500um IAA, the cells were trypsinized, washed twice in fresh media and replated in 6-well plates in fresh media. They were then crosslinked with 1% formaldehyde directly in the 6-well plates at various time points after auxin withdrawal (20, 40, 60, 180, 360, 1080, 1440 minutes).

In order to ensure that our results were not due to the HCT-116 cells stalling in mitosis, we also repeated our Hi-C experiments after cell synchronization and arrest of the cells at the G1/S boundary. First, we added 2mM thymidine to arrest the cells in S-phase and incubated the cells for 12 hours. We then trypsinized and spun down the cells and replated in fresh media, allowing the cells to grow for 12 hours to exit from S-phase. We then added mimosine to a final concentration of 400uM and incubated the cells for 12 hours to arrest the cells at the G1/S boundary. We then replaced media with either complete media + 500uM IAA + 400uM mimosine (treated cells) or complete media + 400uM mimosine (untreated cells) and incubated the cells for 6 hours before processing for downstream experiments.

## METHOD DETAILS

### Microscopy

Live HCT116 RAD21-mAC cells in growth medium without phenol red were added to a chambered coverglass (Lab-Tek #155409) 24 hours prior to imaging and incubated at 37°C, 5% CO2, allowing them to attach to the coverglass. One hour before imaging, the growth medium was replaced with 2 μg/ml of Hoechst 33342 in phosphate-buffered saline (PBS) to visualize nuclei. Time-lapse widefield fluorescence microscopy was performed on a DeltaVision OMX microscope (GE Healthcare) equipped with a 37°C incubation chamber, using a 60⨯ oil immersion objective. Cells were treated with 500μM of IAA immediately before imaging. Images were collected every 10 minutes from 0 to 60 minutes following treatment (DAPI filter at 5%T, 100ms for Hoechst; FITC filter at 100%T, 100ms for mClover), and deconvolved using the built-in SoftWoRx software. The Hoechst images were adjusted in Photoshop by increasing brightness by 92 and contrast by 92 in legacy mode (applied equally to the entire image for all timepoints). The mClover images were adjusted in Photoshop by increasing brightness by 164 and contrast by 123 in legacy mode (applied equally to the entire image for all timepoints). The images were merged in Photoshop using the “screen” function.

### In situ Hi-C

We generated 52 *in situ* Hi-C libraries using the MboI restriction enzyme following the protocol described in (Rao et al., 2014) without modifications. In brief, the *in situ* Hi-C protocol involves crosslinking cells with formaldehyde, permeabilizing them with nuclei intact, digesting DNA with a suitable 4-cutter restriction enzyme, filling the 5’-overhangs while incorporating a biotinylated nucleotide, ligating the resulting blunt-end fragments, shearing the DNA, capturing the biotinylated ligation junctions with streptavidin beads, and analyzing the resulting fragments with paired-end sequencing.

We generated 7 libraries each for our main maps (untreated HCT-116 RAD21-mAC cells and HCT-116 RAD21-mAC cells treated for 6 hours with IAA) comprised of two sets of biological replicates each (three and four technical replicate libraries per biological replicate). In addition, we generated four technical replicate libraries each for untreated and treated HCT-116 RAD21-mAC cells after cell synchronization and arrest. Finally, we generated two technical replicate libraries per time point of our auxin treatment and withdrawal time course. Similar results were obtained with Hi-C libraries from synchronized and arrested cells (Fig. S2I-M), so for all analyses presented in the main text and figures of the manuscript (other than the time course analyses), we utilized our high resolution maps from the unsynchronized cells. Further details about the Hi-C libraries and details about which experiments were used in which figures are provided in Table S1.

### ChIP-Seq

ChIP-Seq for H3K27Ac, H3K4me1, H3K4me3, H3K36me3, H3K27me3, H3K9me3, H4K16Ac, H4K20me3, H3K79me2, and H2.AZ was performed using a native ChIP-Seq protocol. Chromatin from untreated HCT-116 RAD21-mAC cells or cells treated for 6 hours with 500uM IAA was digested with Mnase (Sigma) in digestion buffer (50 mM Tris-HCl, pH7.6, 1 mM CaCl_2_, 0.2% Triton X-100, butyrate 5 mM) for 5’ at 37°C and dialyzed against RIPA buffer for 2hrs at 4°C. Five microgram of respective antibody was incubated with 40 μl of Dynabeads Protein A (or G) for 40 min at room temperature. Antibody-bound beads were added to 500 μl of sonicated or Mnase-digested chromatin, incubated at 4°C overnight, and washed twice with RIPA buffer, twice with RIPA buffer containing 0.3M NaCl, twice with LiCl buffer (0.25 M LiCl, 0.5% Igepal-630, 0.5% sodium deoxycholate), once with TE (pH 8.0) plus 0.2% Triton X-100, and once with TE (pH 8.0). ChIP DNA was purified by phenol-chloroform extraction followed by ethanol precipitation. Libraries were prepped for Illumina sequencing and 50bp single-end reads were sequenced on a HiSeq2000 or 2500 (Illumina).

We performed ChIP-Seq for SMC1 and CTCF following the protocol outlined by the ENCODE consortium (Landt et al., 2012). We also performed ChIP for RAD21 using the protocol from the ENCODE consortium above, 0.292ng/ul of DNA were recovered after immunoprecipitation for untreated cells compared to only undetectable levels of DNA for cells treated with 500uM IAA for 6 hours. These ChIP experiments were not processed for further sequencing.

All ChIP-Seq experiments were processed in parallel with whole cell extract input controls.

### PRO-Seq

To measure changes in transcription resulting from cohesin loss, we performed precision run-on sequencing (PRO-Seq) (Kwak et al., 2013), a variant of global run-on sequencing (GRO-Seq) (Core et al., 2008), using a single biotinylated nucleotide (biotin-11-CTP) as previously described (Engreitz et al., 2016). We made one modifications to the protocol: at the end of each biotin enrichment, we eluted biotinylated RNAs from the streptavidin-coated magnetic beads by heating beads in 25 μl of 20 mM Tris-HCl pH 7.5, 10 mM EDTA, 2% N-lauroylsarcosine at 95°C for 5 minutes, followed by a magnetic-bead nucleic acid purification with 20 μl of MyONE SILANE beads as previously described (Engreitz et al., 2015). During the nuclei preparation step, we processed pairs of RAD21-mAC cells with and without auxin treatment in parallel. In addition, we performed PRO-Seq on HCT-116 CMV-OsTIR1 cells, the parental cell line of RAD21-mAC containing the *OsTIR1* gene integrated at the AAVS1 locus and no mAID tags integrated on any protein. By performing PRO-Seq on CMV-OsTIR1 cells with and without auxin treatment, we could control for transcriptional effects of the auxin treatment itself on HCT-116 cells, as well as any consequences of tagging the RAD21 protein.

## QUANTIFICATION AND STATISTICAL ANALYSIS

### Hi-C Data Processing

All Hi-C libraries were sequenced either on an Illumina NextSeq500 (either 80 or 85bp paired-end reads) or a HiSeqX (150bp paired-end reads). All resulting data was processed using Juicer (Durand et al., 2016; Rao et al., 2014). The data was aligned against the hg19 reference genome. All contact matrices used for further analysis were KR-normalized with Juicer.

Loops were annotated in both untreated and treated maps using HiCCUPS (Durand et al., 2016; Rao et al., 2014). Loops were called at 5kb, 10kb, and 25kb resolutions and merged as described in (Rao et al., 2014). Default parameters as described in (Durand et al., 2016; Rao et al., 2014) were used with the exception that an additional enrichment filter was added. We noted that due to karyotypic abnormalities in the HCT-116 cell line, many rearrangements were annotated in both the untreated and treated maps. Since rearrangements appear as very intense pixels off-diagonal, we removed any peak calls that displayed an observed/expected enrichment of >4.5. Empirically, this max threshold removed peak annotations due to rearrangements; notably, nearly the same number of annotated peaks were removed from the untreated and the treated annotations, 277 and 269 respectively. In the end, we annotated 3,170 loops in our untreated maps and 81 loops in our treated maps.

Domains were annotated in both untreated and treated maps using Arrowhead (Durand et al., 2016; Rao et al., 2014). Domains were called at 5kb and 10kb resolutions using default parameters and merged (retaining the 5kb domain annotation for any pair of domains annotated in both the 5kb and 10kb annotations). We annotated 9,845 domains in our untreated maps and 2,090 domains in our treated maps.

### ChIP-Seq Data Processing

All ChIP-Seq data was aligned to hg19 with BWA (Li and Durbin, 2010), deduplicated using PicardTools, and analyzed with MACS 2.0 (Liu, 2014). All data was normalized against the corresponding input control using the ‘-c’ option of MACS 2.0. ChIP-Seq peaks were called using the ‘callpeak’ function of MACS 2.0 with default parameters. Signal tracks were calculated by using the ‘bdgcmp’ option of MACS 2.0 with the ‘FE’ (fold-enrichment) method. All data for downstream analysis was averaged and extracted using either bwtool or the bigWigAverageOverBed utility from UCSC.

### PRO-Seq Data Processing

For analysis of PRO-Seq data, we aligned 30-bp paired-end reads to the hg19 reference (bowtie2 v2.1.0, (Langmead and Salzberg, 2012)), removed duplicate reads (Picard http://picard.sourceforge.net), and discarded reads with MAPQ < 30 (samtools v.0.1.19 https://github.com/samtools/samtools). We counted reads overlapping RefSeq genes (collapsed by gene symbol to the longest isoform) — this quantification procedure includes signal both at the paused position (near the TSS) as well as in the gene body. We identified genes showing significant differences in transcription with DESeq2 (Love et al., 2014), excluding genes with zero coverage in all samples and calling significance at Benjamini-Hochberg corrected p-value < 0.05.

To determine whether there were global changes in the total amount of transcription (up or down) that would affect the normalization and analysis of these experiments, we included a spike-in control in three of the four PRO-Seq replicates for each of untreated and treated RAD21-mAC and CMV-OsTIR1 cells. Specifically, we added ~500,000 Drosophila S2 cells at the beginning of the protocol, as previously described (Mahat et al., 2016). Upon sequencing of these libraries, we counted the number of spike-in reads by aligning to the Drosophila genome (dmel3) with bowtie2 v2.1.0. We observed similar fractions of reads mapping to the Drosophila spike-in in the matched pairs of degron-expressing and control replicate experiments, indicating that there are not significant global changes in the total amount of transcription upon cohesin loss.

### Random Shuffle Annotations

When performing quantitative analyses on our feature annotations, it was frequently desirable to have a “random control” for the feature annotation in question. We generated such annotations through a random permutation procedure. For one-dimensional features, such as peak loci, we randomly placed the one-dimensional features throughout the genome such that (1) the number of features on any one chromosome stayed the same; (2) the random features did not overlap any gaps in the assembly (i.e. centromeres, telomeres, etc.). Similarly, for two-dimensional features (domains, peaks), we randomly placed the two ends of the features across the genome such that (1) the size distribution of the twodimensional features stayed the same; (2) the number of features on any one chromosome stayed the same; (3) the interval between the ends of the randomized two-dimensional features did not overlap any gaps in the assembly.

### Analysis of CTCF and cohesin binding

In order to confirm the degradation of RAD21 in another way, we performed ChIP for RAD21 in untreated and treated (for 6 hours) HCT-116 RAD21-mAC cells. While we recovered.292 ng/ul of DNA after immunoprecipitation for untreated cells, we did not recover detectable levels of DNA after immunoprecipitation for treated cells, indicating the degradation and lack of chromatin-bound RAD21. The pulled down DNA was not prepared for sequencing.

In order to confirm that degradation of RAD21 resulted in abrogation of full cohesin complex binding to chromatin, we performed ChIP-Seq for SMC1 (see above for experimental details). We visually confirmed that cohesin binding was significantly diminished (see Fig. 1C, Fig. S1A,B). We also analyzed the SMC1 signal in aggregate at the top 20,000 RAD21 peaks called in both replicates of a RAD21 ChIP-Seq experiment performed by ENCODE in wild-type HCT-116 cells. We saw an average 78% reduction in binding strength (mean enrichment = 28.05 in untreated HCT-116 RAD21-mAC cells; mean enrichment = 6.17 in treated cells). These results demonstrate that we were able to quickly abrogate cohesin binding to chromatin to near completion using our auxin-inducible degron system.

**Figure S1.**
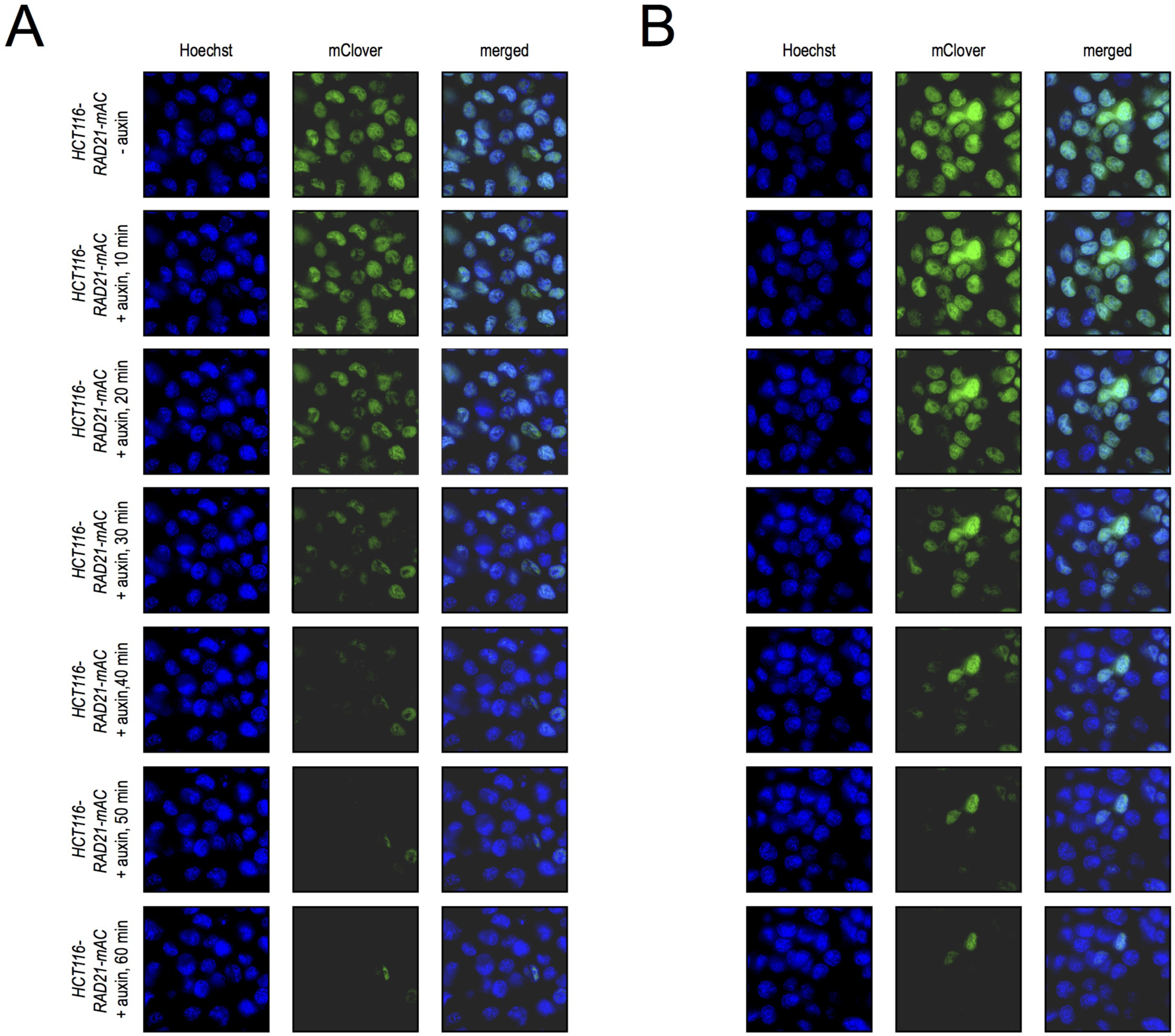
Live cell imaging of HCT-116-RAD21mAC during 1 hour of auxin treatment, Related to Figure 1. (A) Live cell imaging of HCT-116-RAD21-mAC cells after Hoechst 33342 staining to label nuclei. After addition of auxin, nuclear mClover signal corresponding to tagged RAD21 protein rapidly disappears and is nearly completely lost by 1 hour. (B) Another field, treated as above.

We also performed ChIP-Seq for CTCF to establish whether CTCF binding was dependent on cohesin binding or not. While our CTCF ChIP-Seq experiments were noisy, both visual inspection and aggregate analysis of signal at all peaks called in both replicates of a CTCF ChIP-Seq experiment performed by ENCODE in wild-type HCT-116 cells clearly demonstrated that CTCF remained bound after RAD21 degradation. The average enrichment at all CTCF binding sites called by ENCODE was 7.35 in untreated cells and 10.81 in treated cells (The difference in enrichment was likely due to differential quality of the immunoprecipitation rather than true biological differences, but we cannot distinguish between those two possibilities.) This data supports a model in which cohesin is a member of an extrusion complex that slides across DNA, whereas CTCF finds it binding sites independently of the extrusion complex and acts as an oriented brake to halt extrusion complex translocation.

#### Analysis of histone modification patterns

We calculated average signal over 5kb bins across the entire genome and correlated signal between ChIP-Seq experiments before and after auxin treatment for each of CTCF, H3K27me3, H3K9me3, H3K4me1, H3K4me3 and H3K27Ac. All modifications showed high correlations before and after auxin treatment (Spearman’s r = 0.80 [CTCF]; 0.95 [H3K27me3]; 0.95 [H3K9me3]; 0.94 [H3K4me1], 0.70 [H3K4me3]; 0.90 [H3K27Ac]; 0.96 [H2.AZ]; 0.94 [H3K36me3]; 0.96 [H3K79me2]; 0.89 [H4K20me3]; 0.95 [H4K16Ac]).

#### Evaluation of loop and domain formation

We used HiCCUPS to calculate local enrichments on treated maps for the 3, 170 loops we annotated with HiCCUPS in untreated maps. No loop showed at least 1.3-fold enrichment over local backgrounds and <30% FDR q-value. This clearly demonstrates that the vast majority of looping is lost after cohesin degradation. *!*

We identified loop domains as in (Rao et al., 2014), by searching for loop-domain pairs where the peak pixel was within the smaller of 50kb or 0.2 of the length of the domain at the corner of the domain. Using this procedure, we identified 2, 140 loop domains in untreated cells and only 9 in treated cells. Of the 9, 8 were false positives due to rearrangements in HCT-116 cells and one was a false positive due to extensive compartmentalization that was mistakenly annotated as a loop by HiCCUPS. Notably, the high false discovery rate after auxin treatment is due to the very small number of true positives (in this case, no detectable true positives). The false discovery rates of HiCCUPS and Arrowhead before auxin treatment were comparable to the FDRs documented in (Rao et al., 2014). This clearly demonstrates that loop domains are lost after cohesin degradation.

We also assessed the loss of loop domains and loops via aggregate peak analysis (APA). We used default parameters at 10kb resolution, excluding loop domains and loops within 300kb of the diagonal to avoid distance decay effects and extracting a 200kb by 200kb submatrix around every loop domain or loop. In aggregate, the signal from loop domains and loops was clearly and completely lost after auxin treatment: the APA score (fold-enrichment of the peak pixel over the mean value of the 36 pixels in the 6ͯ6 box in the lower left of the aggregate matrix) went from 2.102 to 0.782 for loop domains and 2.095 to 0.797 for all loops. (The APA scores <1 after treatment are expected since random pixels would show an APA score <1 because of the contact probability distance decay.) All visual signs of looping and domain formation were also lost in the aggregate matrices (Fig. S2A, B).

**Figure S2:**
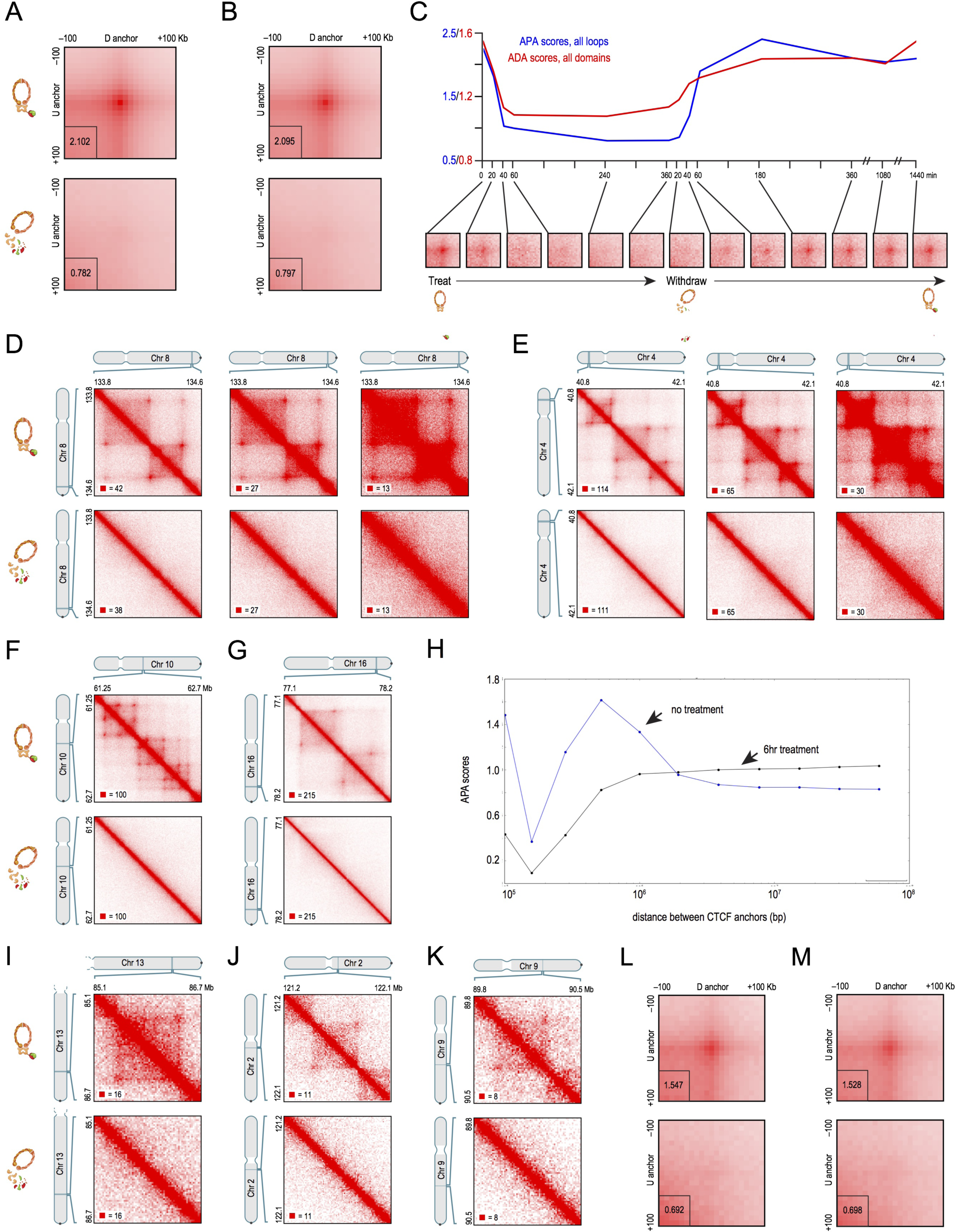
Cohesin degradation eliminates loop domains and the vast majority of loops, Related to Figure 2. (A) APA matrices for all loop domains >300kb long in Hi-C maps for untreated cells (top) versus in Hi-C maps for treated cells (bottom). APA scores are indicated in the lower left corner of each matrix. The color scale for both matrices ranges from 0 (white) to five times the mean of the 6x6 box in the upper right corner of the matrix (red). The APA score after auxin treatment shows complete loss of loop signal (APA score <=1), and no focal enrichment is visible. (B) APA matrices for all loops identified with HiCCUPS with default parameters >300kb long in Hi-C maps for untreated cells (top) versus in Hi-C maps for treated cells (bottom). The color scale for both matrices ranges from 0 (white) to five times the mean of the 6x6 box in the upper right corner of the matrix (red). The APA score after auxin treatment shows complete loss of loop signal (APA score <=1), and no focal enrichment is visible. (C) APA was used to measure the aggregate strength of the links associated with all loops in low-resolution Hi-C contact maps generated across a time course of auxin treatment and withdrawal. APA scores are shown on top; values greater than 1 indicate the presence of loops. APA plots for each time point are shown on the bottom; the strength of looping is indicated by the extent of focal enrichment at the center of the plot. Loops are rapidly lost as cohesin is degraded, and quickly restored when auxin is withdrawn. ADA was used to measure the aggregate gradient across domain boundaries for all domains annotated in untreated cells in low-resolution Hi-C contact maps generated across a time course of auxin treatment and withdrawal. Domain signal is rapidly lost after auxin treatment, but does not completely disappear (reach 1) consistent with the presence of compartment domains after cohesin degradation. (D) An example region from Figure 2A shown at different color scales: contact maps from untreated cells on top and maps from auxin treated cells on the bottom. The lack of visible loop domain structure is not a result of color scale choice; there is no residual loop domain structure. (E) Another example region from Figure 2A shown at different color scales; again there is no residual loop domain structure. (F) An additional example (chr10: 61.25-62.7 Mb) of complete loss of loop domains after auxin treatment and degradation of cohesin. (G) An additional example (chr16: 77.1-78.2 Mb) of complete loss of loop domains after auxin treatment and degradation of cohesin (H) APA scores vs. distance for pairs of convergently oriented CTCF/cohesin-associated loop anchors separated by a given distances. In untreated maps, positive APA scores can be seen for convergently oriented pairs of CTCF/cohesin-associated loop anchors up to distances less than a few megabases, but rapidly drops off at longer distances. In treated maps, positive APA scores are not seen at any distance for convergently oriented pairs of CTCF/cohesin-associated loop anchors. (I) To assure that the disappearance of loop domains after cohesin degradation did not arise as a result of cell cycle abnormalities, we performed Hi-C on cells that were synchronized and arrested at the G1/S boundary before and during auxin treatment. Here, we show an example of a loop domain (chr13: 85.1-86.7 Mb) that is present in our maps from G1-arrested cells and lost after auxin treatment. (J) Another example of a loop domain (chr2: 121.2-122.1 Mb) that is present in our maps from G1-arrested cells and lost after auxin treatment. (K) Another example of a loop domain (chr9: 89.8-90.5 Mb) that is present in our maps from G1-arrested cells and lost after auxin treatment. (L) APA matrices for all loop domains >300kb long (called in our untreated, unsynchronized maps) in Hi-C maps for untreated G1-arrested cells (top) versus in Hi-C maps for treated G1-arrested cells (bottom). The color scale for both matrices ranges from 0 (white) to five times the mean of the 6ͯ6 box in the upper right corner of the matrix (red). The APA score after auxin treatment shows complete loss of loop signal (APA score <=1), and no focal enrichment is visible. (M) APA matrices for all loops >300kb long (called in our untreated, unsynchronized maps) in Hi-C maps for untreated G1-arrested cells (top) versus in Hi-C maps for treated G1-arrested cells (bottom). The color scale for both matrices ranges from 0 (white) to five times the mean of the 6x6 box in the upper right corner of the matrix (red). The APA score after auxin treatment shows complete loss of loop signal (APA score <=1), and no focal enrichment is visible.

In order to assess the dynamics of loop and domain formation we used APA and aggregate domain analysis (ADA) to assess loop, loop domain, and domain strength across a time course of auxin treatment and withdrawal.

ADA uses the same principle of aggregating submatrices across a feature list, but instead of calculating a score representing the focal enrichment of a peak pixel against pixels to its lower left (the APA score), we calculate a score representing the enrichment of contacts just inside the domain boundaries over the contacts just outside the boundary, i.e. a gradient across the boundary. More specifically, we compare the average contacts in the pixels [i+3, j-13:j-3], [i+4,j-12:j-2], [i+5,j-11:j-1], [i+1:i+11,j-5], [i+2:i+12,j-4], [i+3:i+13,j-3] (the inside domain pixels) to the pixels [i-5,j-21:j-11], [i-4,j-20:j-10], [i-3,j-19:j-9], [i+11:i+21,j+5], [i+10:i+20,j+4], [i+9:i+19,j+3] (the outside domain pixels) where (i,j) is the center of the aggregate matrix (i.e. the corner of the domains). Here, we extracted a 200kb by 200kb matrix at 5kb resolution around every domain corner.

For APA on the time course experiments, as with the APA on our deep maps, we used default parameters at 10kb resolution. In Fig. 2B, we show the APA scores for all loop domains greater than 300kb in size. The APA scores demonstrate that after cohesin is degraded, loop domains are completely lost between 40-60 minutes after treatment. From our own imaging (Fig. S1) and imaging performed in Natsume et al. (2016), we know that the half life of cohesin after auxin treatment is about 20 minutes. Thus, loop domains are lost within minutes of cohesin degradation, indicating that cohesin is required for active maintenance of loop domain structures, not just establishment. After withdrawal of auxin, loop domains form similarly quickly, with strong loop domain signal by 60 minutes after withdrawal. This time includes the time for cohesin levels to recover and thus likely represents a very conservative upper bound on the time required for loop domain formation. Additionally, it indicates that loop domain structures are dynamically maintained during interphase.

In Fig. S2C, we show similar results for all loops greater than 300kb in size called in untreated cells. Additionally, we show ADA scores for all contact domains larger than 300kb in size called in untreated cells. While the ADA scores for all domains show a sharp decline in the first 60 minutes after auxin treatment, they plateau above 1, indicating some residual domain signal from al domains. Since, as we noted in (Rao et al., 2014), not all domains are loop domains, this suggest residual retention of non-loop domains. As we noted in (Rao et al., 2014), non-loop domains are usually created by compartment intervals. The retention of compartment domains is discussed below.

#### Analysis of previous cohesin-depletion Hi-C data sets

Previous Hi-C studies after cohesin or CTCF depletion showed limited effects, with both contact domains and compartments present after depletion (Seitan et al., 2013; Sofueva et al., 2013; Zuin et al., 2014). However, in these studies, the authors performed low resolution Hi-C experiments, raising the possibility that either (i) the authors could not resolve the difference between loop domains (which disappear after cohesin loss) and compartment domains (which remain) due to resolution issues, or (ii) incomplete depletion of cohesin or CTCF led to modest phenotypes. The authors in all three studies acknowledge the possibility that the limited effects they see may have been due to incomplete depletions.

To test this hypothesis, we re-analyzed data from these three studies. We downloaded the raw fastqs for all Hi-C experiments performed in the studies and processed them with Juicer (in exactly the same way that we processed all the Hi-C data generated for this study. Although the experiments did not have sufficient resolution to visualize individual loops, we looked for the statistical signal of loop enrichment in aggregate using APA (Durand et al., 2016; Rao et al., 2014). For the mouse data sets generated in Seitan et al. and Sofueva et al., we used a loop list we had previously generated in CH12-LX mouse lymphoblast cells (Rao et al., 2014) with the added filter that we removed loops with >4.5 enrichment over local background in order to stay consistent with the methods used in this study (see above). For the human data sets generated in Zuin et al., we used the loop list of 3,170 loops in untreated HCT-116 RAD21-mAC cells described above. We observed positive APA scores (>1) and visible focal enrichment in all experiments generated in previous studies, before and after cohesin or CTCF depletion (Fig. S3). By contrast, our maps after auxin treatment show complete loss of APA signal and no visible focal enrichment, even when APA is performed on low resolution data sets (Fig. 2B). In previous studies, the APA score was weaker after cohesin or CTCF depletion but still clearly visible and notably, positive APA signal was seen in every replicate experiment performed in previous studies. Taken together, this suggests that a major confound of previous studies was the incomplete depletion of cohesin or CTCF, and along with the limited resolution of the Hi-C experiments, likely explains the limited effects seen.

**Figure S3:**
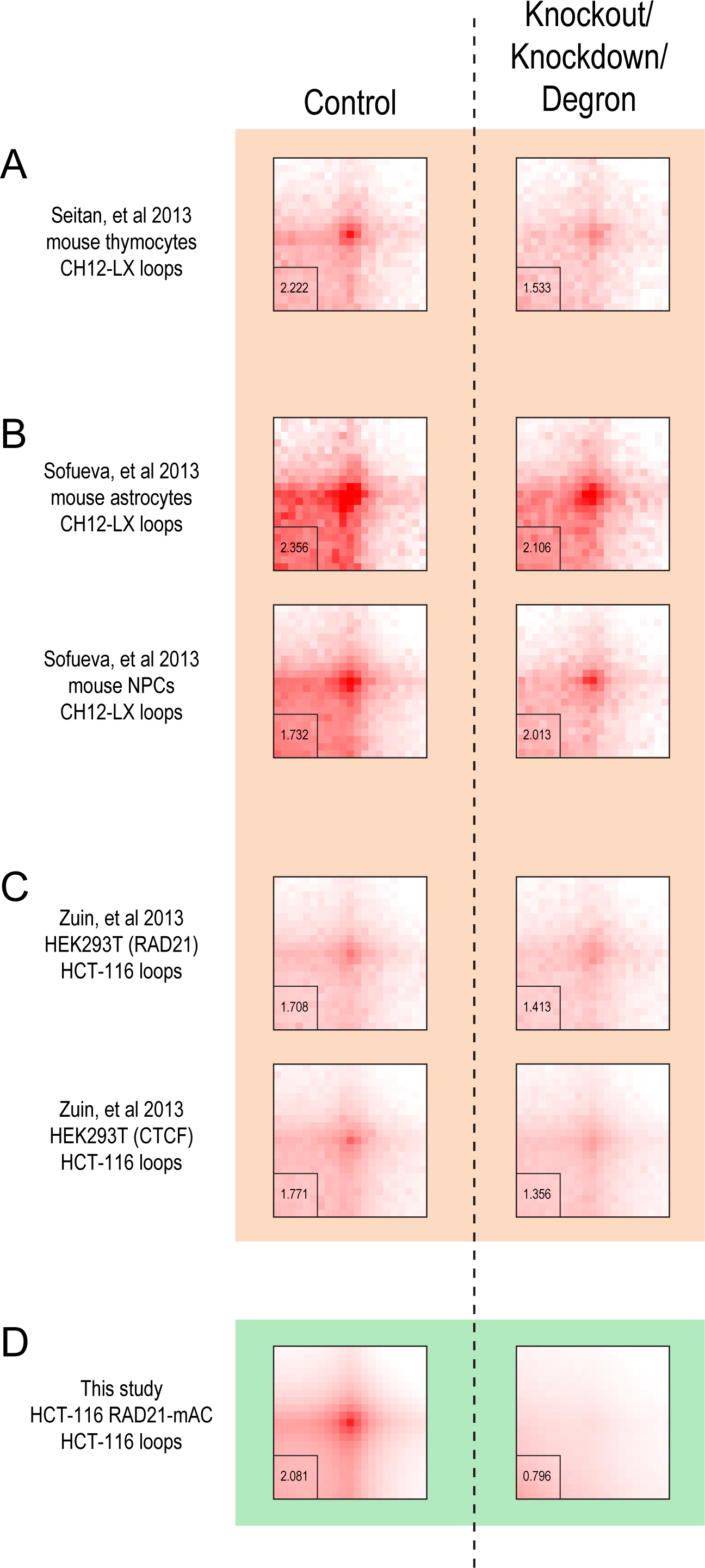
Previous low-resolution Hi-C studies of cohesin and CTCF depletion show signs of loop domains suggesting incomplete depletion, Related to Figure 2. (A) We re-analyzed the Hi-C data from (Seitan et al., 2013) and performed APA on their maps from mouse thymocytes before cohesin deletion (left) and after (right) using a loop list we generated using HiCCUPS in CH12-LX mouse lymphoblasts (Rao et al., 2014). A positive APA score (1.533, indicating ~1.5 fold enrichment of the peak pixel over the pixels to its lower left) is seen even after cohesin deletion suggesting incomplete deletion of RAD21. The color scale for all APA matrices in this figure ranges from the mean of the 6x6 box in the upper right corner (white) to five times the mean of the 6ͯ6 box in the upper right corner of the matrix (red). (B) We re-analyzed the Hi-C data from (Sofueva et al., 2013) and performed APA on their maps from mouse astrocytes before cohesin deletion (top left) and after (top right) as well as on their maps from mouse NPCs before cohesin deletion (bottom left) and after (bottom right) using a loop list we generated using HiCCUPS in CH12-LX mouse lymphoblasts (Rao et al., 2014). A positive APA score (2.106 and 2.013 respectively, indicating ~2 fold enrichment of the peak pixel over the pixels to its lower left) is seen even after cohesin deletion suggesting incomplete deletion of RAD21. (C) We re-analyzed the Hi-C data from (Zuin et al., 2014) and performed APA on their maps from HEK293T cells before cohesin depletion (top left) and after (top right) as well as on their maps from HEK293T cells with a control siRNA (bottom left) and with an siRNA targeting CTCF (bottom right) using the loop list generated with HiCCUPS in untreated HCT-116 RAD21-mAC cells in this study. A positive APA score (1.413 and 1.356 respectively, indicating ~1.4 fold enrichment of the peak pixel over the pixels to its lower left) is seen even after cohesin or CTCF depletion suggesting incomplete depletion. (D) By contrast, APA on our maps from untreated HCT-116 RAD21-mAC cells and treated HCT-116 RAD21-mAC cells with the untreated RAD21-mAC loop list shows that after complete cohesin depletion, focal enrichment is completely lost in the APA matrix and the APA score is <=1.

#### Evaluation of genome compartmentalization

The most common method used for classifying Hi-C patterns is the principal component (PC) approach, which we introduced in (Lieberman-Aiden et al., 2009). In this approach, each intrachromosomal contact matrix is converted to an observed/expected matrix, and the first principal component of this matrix is used to bifurcate the data into two clusters. We showed in (Rao et al., 2014) that this method does not capture compartment structure accurately at high resolutions; however it is useful for comparing gross compartmentalization patterns.

We first calculated the first three principal components of the 25-kb resolution observed/expected matrix for each chromosome (constructed using Juicer as described in *1,4,13)* using scikit learn’s RandomizedPCA function. We chose the principal component most correlated with GC content and assigned sign such that the vector was positively correlated with the GC content vector. We then calculated the correlation of the eigenvector for each chromosome between untreated and treated maps. The mean correlation was 0.968.

To identify transitions in compartment state at higher resolution, we used a combination of techniques. First, we calculated an edge score using an algorithm similar to Canny edge detection. For every 25kb locus in the genome, we looked at the corresponding column of the 25kb log2(observed/expected) matrix. For every pixel (i,j) in column j, we calculated a gradient = [i,j:j+3] - [i,j-4,j-1]. We then searched for stretches of at least 7 pixels in the column with a gradient x such that abs(x) was greater than 0.5. We then extended the edges by including pixels adjacent to an edge that had a gradient of at least 0.3. Finally, we summed the number of pixels in a column belonging to an edge to calculate the edge score for a locus. We then called local peaks in this track which could correspond to compartment state transitions since a compartment state transition at locus I will create an edge between locus i-1 and i.

Since loci in the same compartment will exhibit the same rises and falls in contact probability as one slides along the genome, we reasoned that adjacent pixels should exhibit high correlations of the derivative of their contact patterns and low correlations could indicate a compartment state transition. As described in Section V.a.3 of Rao et al., 2014, this is akin to measures in finance that correlate returns of prices to identify similarities between stocks. To calculate this sliding derivative correlation score, we calculated the gradient in the log2(observed/expected) matrix over every boundary called in our edge score track. More specifically, for every locus i, and all boundaries j in boundary set J that were within 15Mb of i, we calculated the difference of mean([j:j+5,i]) and mean([j-6:j-1,i]). We then calculated the Spearman correlation coefficient of these two vectors (one vector for the gradients at all boundaries j in J for the pixels upstream of i, and one vector for the pixels downstream of i). We excluded the derivative signal at pixels not located at compartment state transitions as defined by the edge score to reduce noise, reasoning that pixels inside compartment intervals were unlikely to contribute meaningful rises/falls in contact probability. Similarly, we only included pixels within 15mb of i to reduce the noise arising from sparsity far off the diagonal.

We then identified compartment boundaries by calling local peaks in the edge score track and local valleys in the sliding derivative correlation score track and merging the two peak call lists. We identified 4,325 boundaries in untreated cells for a median compartment size of 425 kb and 4,424 boundaries in treated cells for a median compartment size of 475 kb. These are very likely conservative upper bounds on the true median compartment size, since we utilized stringent peak calling and compartment structure can be difficult to detect in maps that are not extremely high resolution (Rao et al., 2014).

To assess the presence and strength of contact domains after auxin treatment, we used the Arrowhead algorithm (Durand et al., 2016; Rao et al., 2014). The Arrowhead algorithm calculates a corner score for every pixel, where higher corner score values represent a higher likelihood that a pixel is at the corner of a domain (see Section IV.a.3 of the Extended Experimental Procedures of Rao et al., 2014). For our list of 9,845 contact domains identified by the Arrowhead algorithm in untreated RAD21-mAC cells, we compared the corner scores of the contact domains to the corner scores of random pixels with an identical chromosome and length distribution. The median corner score in untreated cells for all domains called in untreated cells was the 97th percentile of random corner scores. We then calculated the corner scores in treated cells for the list of contact domains annotated in untreated cells, as well as the corner scores for the random control. Here, the median corner score for annotated contact domains was only the 86th percentile of random corner scores. (Notably, the distributions of scores for random pixels did not change, see Fig. 3B.) This indicates that contact domains were significantly weakened after auxin treatment. However, there was still some residual signal.

Since we knew that loop domains were completely eliminated from our previous analyses and that compartment structure remained after treatment, we reasoned that the residual signal was arising from retained compartment domains (contact domains whose boundaries overlap compartment interval boundaries). To test this, we identified 974 contact domains whose boundaries overlapped a compartment interval boundary (within 25 kb), i.e. compartment domains. Additionally, we identified 410 contact domains whose boundaries were not within 100 kb of a compartment boundary even after using a relaxed threshold for identifying compartment boundaries, i.e. a high confidence set of non-compartment domains. We then analyzed the corner scores for each of these sets of domains separately in treated cells and found that while the median score for compartment domains was 89^th^ percentile of the random corner scores, the median score for non-compartment domains was only 72nd percentile of the random corner scores. This indicates that the residual signal stems from retained compartment domains. Thus, while loop domains are completely eliminated, contact domain structure arising from genome compartmentalization is still present after auxin treatment, although the domains are weaker than those found in untreated cells.

It is commonly thought in the literature that contact domains and compartment intervals form a hierarchy, with compartment intervals often being subdivided into multiple contact domains, but each contact domain belonging to only one compartment interval. Having determined that loop domains and compartmentalization formed via independent mechanisms, we wondered whether loop domains and compartment intervals shared characteristic hierarchical relationships or whether they truly formed independently in the genome.

To assess whether compartment boundaries could be spanned by loop domains, we intersected our loop domain annotation and our compartment boundary annotation. Specifically, we identified compartment boundaries in our treated maps that were contained within a loop domain called in untreated cells and >100kb away from either loop anchor (obviously this excludes loop domains smaller than 200kb from the analysis). We identified 349 such boundaries. Visual examination also confirmed that these boundaries were true compartment state transitions lying inside loop domains (Fig. 3C,D).

Note that this is a lower bound on the number of compartment boundaries spanned by loop domains, as we used stringent distances from loop anchors to reduce false positives and our compartment boundary annotation has false negatives as well. This demonstrates that there is no true hierarchy between compartmentalization and loop domain formation, contrary to what has been suggested in the literature.

We wondered what happen to compartment strength at these boundaries when loop domains were eliminated. To analyze this, we calculated the average sliding derivative correlation score (see above) for the 1Mb intervals centered on the 349 compartment boundaries contained within loop domains before and after auxin treatment. We observed that the boundaries contained within loop domains showed a strong increased in compartment strength (larger dip in the sliding correlation score) after the elimination of loop domains: 0.10 decrease in the sliding correlation score in untreated cells vs. 0.31 in treated cells. In contrast, when we identified 389 compartment boundaries in treated cells that were positioned at loop domain anchors annotated in untreated cells (within 25kb), we found that there a much more modest increase in compartment strength after treatment: 0.35 decrease in the sliding correlation score in untreated cells vs. 0.53 in treated cells (Fig. 3E). This indicates that cohesin facilitates mixing of distinct compartment states and causes decreases in compartmentalization unless it is halted at the compartment boundary.

The results were similar when we examined compartment boundaries inside all loops: we identified 593 compartment boundaries in treated cells that were spanned by loops and at least 100kb away from either loop anchor, and we identified 503 compartment boundaries in treated cells that were positioned at loop anchors. We saw an 0.11 decrease in the sliding correlation score in untreated cells vs. 0.37 decrease in treated cells for compartment boundaries spanned by loops, and an 0.38 decrease in the sliding correlation score in untreated cells versus an 0.54 decrease in treated cells for compartment boundaries at loop anchors (Fig S3B).

To assess whether the changes in compartmentalization seen after treatment corresponded to epigenetic activity, we performed a similar analysis except instead of calling compartment boundaries, we identified transitions in broad histone modification state for H3K27Ac and H3K27me3. Since histone modifications have been shown to very closely correlate with compartmentalization (Lieberman-Aiden et al., 2009; Rao et al., 2014; Sexton et al., 2012), we reasoned that changes in histone modification within loop domains and loops should show greater changes in compartmentalization to better match the histone modification pattern compared to changes in histone modification status at loop anchors. We identified changes in H3K27Ac status by creating a 25kb binary track that was either 0 if the enrichment was less than 0.35 or 1 if the enrichment was greater than 0.35. We then calculated the absolute value of a smoothed gradient (using the kernel [1 1 1 -1 -1 -1]) and called local peaks to identify changes in histone modification status. We identified 264 H3K27Ac transitions spanned by loop domains (same definition as above) and 307 H3K27Ac transitions positioned at loop domain anchors. The H3K27Ac signal in the 1Mb intervals around these transitions did not change after auxin treatment (Fig. 3F). However, while there was very little change in the compartmentalization strength at transitions at loop domain boundaries (0.41 dip in sliding correlation in untreated vs. 0.49 in treated), there was a dramatic increase in compartmentalization strength at transitions spanned by loop domains (0.02 dip in sliding correlation in untreated vs. 0.19 in treated). This indicates that removal of loop domains by cohesin loss leads to genome compartmentalization that more closely matches histone modification patterns.

Similar results were seen for H3K27Ac transitions spanned by all loops: we identified 426 H3K27Ac transitions in untreated cells that were spanned by loops and at least 100kb away from either loop anchor, and we identified 381 H3K27Ac transitions in untreated cells that were positioned at loop anchors. The H3K27Ac signal in the 1Mb intervals around these transitions did not change after auxin treatment (Fig. 3F). We saw an 0.41 decrease in the sliding correlation score in untreated cells vs. 0.50 decrease in treated cells for H3K27Ac transitions spanned by loops, and an 0.10 decrease in the sliding correlation score in untreated cells versus an 0.26 decrease in treated cells for H3K27Ac transitions at loop anchors (Fig S4B).

**Figure S4:**
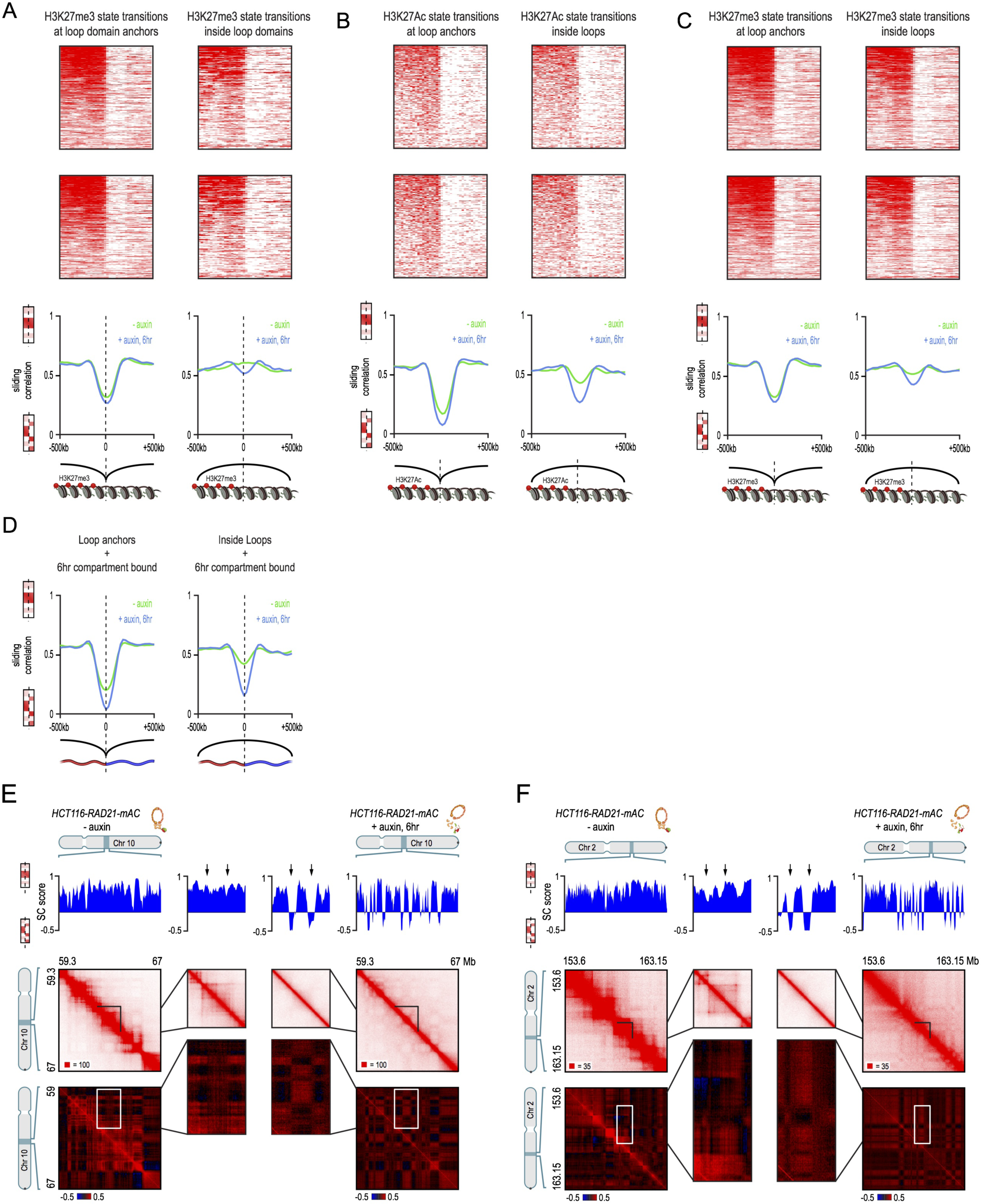
Cohesin degradation results in genome compartmentalization that better matches histone modification patterns, Related to Figure 3. (A) Sliding correlation scores before and after auxin treatment for H3K27me3 domain boundaries in untreated cells which either coincide with loop domain anchors (left) or are located in the interior of a loop domain (right). H3K27me3 histone modification patterns do not change after loss of cohesin (top and middle). For H3K27me3 boundaries that lie in the interior of a loop domain in untreated cells, the difference in long-range contact pattern on opposite sides of the boundary increases greatly after cohesin treatment. This indicates that loop domains facilitate mixing of chromatin with different histone modifications. (B) Sliding correlation scores before and after auxin treatment for H3K27Ac domain boundaries in untreated cells which either coincide with loop anchors (left) or are located in the interior of a loop (right). For H3K27Ac boundaries that lie in the interior of a loop in untreated cells, the difference in long-range contact pattern on opposite sides of the boundary increases greatly after cohesin treatment. This indicates that loops facilitate mixing of chromatin with different histone modifications. (C) Sliding correlation scores before and after auxin treatment for H3K27me3 domain boundaries in untreated cells which either coincide with loop anchors (left) or are located in the interior of a loop (right). For H3K27me3 boundaries that lie in the interior of a loop in untreated cells, the difference in long-range contact pattern on opposite sides of the boundary increases greatly after cohesin treatment. This indicates that loops facilitate mixing of chromatin with different histone modifications. (D) Sliding correlation scores before and after auxin treatment for compartment boundaries which either coincide with loop anchors (left) or are located in the interior of a loop (right). For compartment boundaries that lie in the interior of a loop in untreated cells, the difference in long-range contact pattern on opposite sides of the boundary increases greatly after cohesin treatment. (E,F) Examples (E: chr10:59.3-67Mb and D: chr2:153.6-163.15Mb) showing that the loss of cohesin-associated loops after auxin treatment results in increased fine-scale compartmentalization. Top: Sliding correlation scores; valleys imply strong differences in long-range contact pattern observed at a locus as compared to neighboring loci, indicating a change in compartment (see Methods). Middle: Observed contact matrices. Bottom: Pearson’s correlation maps for the local region shown (see Methods). Deeper valleys in the sliding correlation score and increased plaid patterning in the observed and Pearson’s correlation maps indicate strengthened fine-scale compartment interactions after auxin treatment. Blowouts: loss of a loop domain results in strengthening of a compartment boundary spanned by the loop. Blown-out regions are indicated on zoomed out maps for both the observed (black upper triangle) and Pearson’s correlation maps (white rectangle). Observed and Pearson’s correlation maps are both shown at 25kb resolution for the zoomed out matrices and 10kb and 25kb resolution respectively for the blown-out matrices.

We also performed this analysis for H3K27me3. We calculated the gradient at every 25kb locus i the genome by taking the absolute value of the difference between the summed log2 fold-enrichment for pixels i-8 to i-1 and the summed log2 fold-enrichment for pixels i+1 to i+8. We called local peaks on this gradient track to identify loci where the broad H3K27me3 modification status changed. We identified 209 H3K27me3 transitions spanned by loop domains (same definition as above) and 384 H3K27me3 transitions positioned at loop domain anchors. The H3K27me3 signal in the 1Mb intervals around these transitions did not change after auxin treatment (Fig. S4A). However, while there was very little change in the compartmentalization strength at transitions at loop domain boundaries (0.29 dip in sliding correlation in untreated vs. 0.33 in treated), there was a stronger increase in compartmentalization strength at transitions spanned by loop domains (0.01 increase in sliding correlation in untreated vs. 0.03 dip in treated).

Similar results were seen for H3K27me3 transitions spanned by all loops: we identified 391 H3K27me3 transitions in untreated cells that were spanned by loops and at least 100kb away from either loop anchor, and we identified 469 H3K27me3 transitions in untreated cells that were positioned at loop anchors. The H3K27me3 signal in the 1Mb intervals around these transitions did not change after auxin treatment (Fig. S4C). We saw an 0.27 decrease in the sliding correlation score in untreated cells vs. 0.31 decrease in treated cells for H3K27me3 transitions spanned by loops, and an 0.03 decrease in the sliding correlation score in untreated cells versus an 0.12 decrease in treated cells for H3K27me3 transitions at loop anchors (Fig S4C).

Taken together, these results suggest that cohesin facilitates mixing of chromatin with different histone modification states and loss of cohesin leads to better correspondence of genome compartmentalization with histone modification patterns and gene activity.

#### Annotation and analysis of cohesin-independent links

We first annotated loops in our maps for auxin-treated RAD21-mAC cells using default HiCCUPS parameters for 5,10, and 25kb resolutions (Durand et al., 2016; Rao et al., 2014) with the additional requirement that the peak pixel show less than 4.5-fold enrichment over local expecteds (in order to remove as many false positives as possible due to rearrangements and assembly issues, see above). Using this procedure, we annotated 81 loops in treated RAD21-mAC cells. When we visually examined these loops, we found that 66 were false positives, with 55 of the false positives due to assembly issues, issues with repetitive elements or structural rearrangements. The false discovery rate for HiCCUPS is much higher in treated cells because the number of true positives is dramatically lower. As mentioned above, the false discovery rate in untreated cells was comparable to the rates described previously in (Rao et al., 2014); in fact, as one might expect false positives to arise from artifacts in the data that are independent of cohesin-mediated looping, the reduction by nearly 98% of numbers of loops called by HiCCUPS after auxin treatment is a powerful proof of its accuracy. When we examined the 15 true positive loops annotated by HiCCUPS, we found that they had a dramatically different distance distribution than cohesin-associated loops: where the median size of a cohesin-associated loop was 275kb, the median size of these 15 loops was 1.75Mb. We also noticed that the anchors involved in these 15 loops were often forming long-range loops at distances of tens of megabases and hundreds of megabases. We reasoned that HiCCUPS using default parameters for loop detection was missing many of these extremely long-range loops because of the extra stringency of the HiCCUPS lambda chunking procedure for multiple hypothesis testing for pixels with low counts (i.e. pixels far off the diagonal). To call more of these long-range loops, we decided to modify the HiCCUPS parameters similar to make the parameters more similar to those used to identify the extremely long-range “superloops” on the inactive X chromosome (Rao et al., 2014; Darrow et al., 2016).

We decided to annotate loops in auxin-treated RAD21-mAC cells with the parameters used in to annotate superloops on the inactive X chromosome (which also form between loci tens to hundreds of megabases apart). More specifically, we annotated loops by running HiCCUPS at 50 and 100kb resolutions with the following parameters: p = 2,1; w = 4,2; fdr = 10%, 10%. We additionally filtered loops that were within 5 Mb of the diagonal, had less than a 2-fold observed/expected for any of the local expected, and had fewer than 3 pixels clustered into the peak pixels (see section VI.a.5 of Rao et al., 2014). This annotation yielded 88 loops. After visual examination, we found that 46 of these loops corresponded to true positives while the other 42 were false positives (22 were due to issues with repetitive regions and 15 were due to other forms of structure in the contact map, for instance interactions between broad compartment intervals). Combining these 46 loops with the 15 loops annotated with high resolution HiCCUPS, we obtained a final curated list of 61 intrachromosomal cohesin-independent loops.

We first identified the loop anchors contributing to the cohesin-independent loops. We merged all adjacent loci involved in one of the 61 loops annotated above. We then expanded all loop anchor loci to be 100kb in size, yielding a list of 64 loop anchor loci.

To assess the presence and orientation of CTCF at loop anchor loci for both cohesin-associated and cohesin-independent loop anchors, we followed the procedure exactly from section VI.e.7 of (Rao et al., 2014). In order to use comparable loop anchor sizes, we collapsed each 100kb cohesin-independent loop anchor to the 15kb interval in the center of the 100kb interval. We found that while 90% of cohesin-associated loop anchors were associated with CTCF binding, only 20% of cohesin-independent loop anchors were associated with CTCF binding. More over, while 95% of unique CTCF motifs in cohesin-associated loop anchors pointed towards the interior of the loop (consistent with the convergent rule), the unique CTCF motifs in cohesin-independent loops did not exhibit any such bias (56% pointing towards the interior of the loop) (Fig. 4B). This strongly suggests that cohesin-independent loops form via a mechanism other than extrusion.

To analyze enrichment of proteins bound at cohesin-independent loop anchors, we reproduced the analysis from section VI.e.7 of (Rao et al., 2014), using the 100kb loop anchors and comparing to the average of 100 randomly shuffled loop anchor lists (see the section on Random Shuffle controls above). We downloaded peak calls for 36 DNA-binding proteins or histone modifications in HCT-116 cells from ENCODE (ENCODE Consortium, 2012). We also utilized an annotation of stitched and ranked (by H3K27Ac enrichment) superenhancers and enhancers from (Hnisz et al., 2013). For each of the proteins or histone modifications, we calculated the percentage of loop anchors that overlap the feature as well as the enrichment over the percentage of random anchors overlapping the feature. We found that strong H3K27Ac sites and superenhancers (especially the strongest 100 superenhancers) were very strongly enriched at cohesin-independent loop anchors (Fig. 4F).

We also performed the analyses listed above on automated lists of cohesin-independent loops without any manual curation. We found that the results showing a lack of CTCF binding at cohesin-independent loop anchors and a lack of CTCF orientation preference were similar (Fig. S5A). We also found that superenhancers were strongly enriched at loop anchors generated from the 88 loop list automatedly called with low resolution HiCCUPS; the top 100 superenhancers were 47-fold enriched (present at 30/115 loop anchors). See figure S5C. This indicates that our results were not biased by our use of a manually curated loop list.

**Figure S5:**
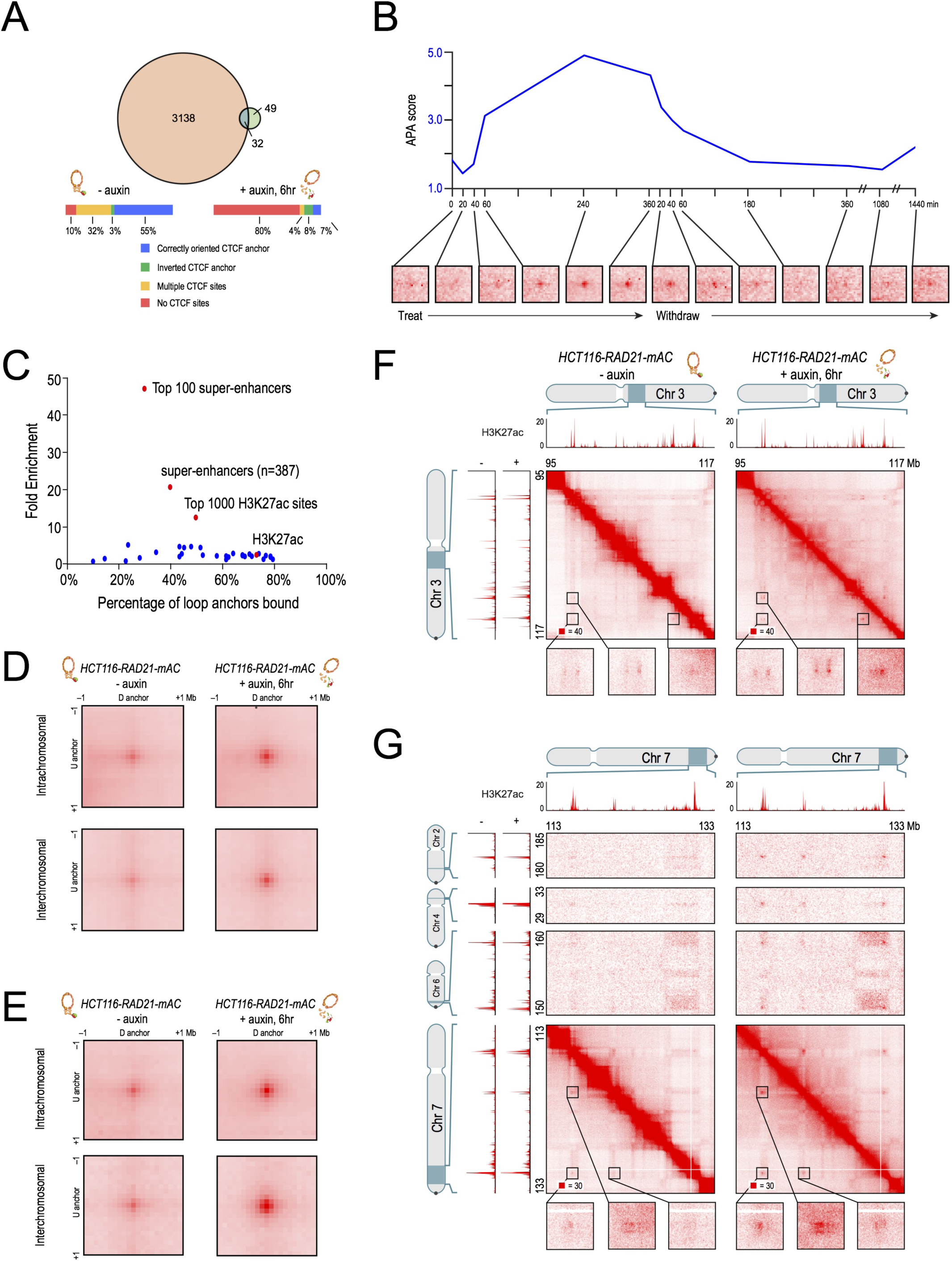
Co-localization of superenhancers after cohesin loss, Related to Figure 4. (A) Venn diagram of loops called in untreated cells with default HiCCUPS parameters with loops called in treated cells with default parameters. The vast majority of loops are lost (>97%) but a small number of “cohesin-independent” loops remain. Loops annotated in treated cells do not frequently bind CTCF and show no CTCF orientation bias. (B) APA for intrachromosomal (blue) cohesin-independent links across a time course of auxin treatment and withdrawal using an automatedly generated list by pairing all superenhancers overlapping cohesin-independent link anchors returned by low-resolution HiCCUPS. APA scores are shown on top and APA plots for each time point are shown on the bottom. Cohesin-independent links are rapidly strengthened as cohesin is degraded and weaken as cohesin is restored. (C) Percent of cohesin-independent loop anchors bound versus fold enrichment for 36 DNA-binding proteins and histone modifications. Same analysis as Fig. 4F but using a loop anchor list generated by running HiCCUPS at 50 and 100kb resolution without any manual curation. Superenhancers are still highly enriched at cohesin-independent loop anchors, validating that the result does not stem from hand curation bias. (D) APA for intrachromosomal (top) and interchromosomal (bottom) cohesin-independent links in our untreated maps (left) and our treated maps (right) using an automatedly generated list by pairing all superenhancers overlapping cohesin-independent link anchors returned by low-resolution HiCCUPS (same list as Fig. S5B and S5C). Cohesin-independent links are significantly strengthened as cohesin is degraded (Intrachromosomal APA scores: 1.69 (untreated) vs. 2.75 (treated); interchromosomal APA scores: 2.29 (untreated) vs. 3.64 (treated)). (E) APA for intrachromosomal (top) and interchromosomal (bottom) cohesin-independent links in our untreated maps (left) and our treated maps (right) using our manually curated HiCCUPS lists of 61 intra- and 203 interchromosomal links. Cohesin-independent links are significantly strengthened as cohesin is degraded (Intrachromosomal APA scores: 2.32 (untreated) vs. 4.02 (treated); interchromosomal APA scores: 3.02 (untreated) vs. 7.02 (treated)). (F) Examples of a network of intrachromosomal cohesin-independent links between superenhancers on chr3. H3K27 acetylation does not change with auxin treatment, but cohesin-independent links are significantly strengthened upon treatment. (G) Examples of a network of intra- and interchromosomal cohesin-independent links between superenhancers on chr7, chr6, chr4, and chr2. H3K27 acetylation does not change with auxin treatment, but cohesin-independent links are significantly strengthened upon treatment.

We noticed that our 64 cohesin-independent loop anchors determined from our 61 loop intrachromosomal list often formed focal interchromosomal links between pairs of loop anchors and that there were large cliques of interactions between anchors (Fig. 4A,B,E). This is in stark contrast to cohesin-associated loop anchors, which show no such enrichment for extremely long Intrachromosomal interactions or interchromosomal interactions, either when examined individually or in aggregate via APA (Fig S2H). This strongly suggests that cohesin-independent loops and links form via a mechanism other than extrusion, since extrusion cannot occur on two topologically distinct molecules‥

To annotate these interchromosomal links between pairs of cohesin-independent loop anchors, we used HiCCUPS to calculate local enrichments at 100kb resolution for all possible interchromosomal pairs of cohesin-independent loop anchors. We then identified enriched focal interchromosomal interactions by filtering for links that were enriched at least 5.5-fold over local background (empirically chosen to ensure a <10% false discovery rate). Using this procedure, we identified 203 interchromosomal cohesin-independent links. This likely underestimates the true number of interchromosomal cohesin-independent links, as evidenced by Fig. 4B.

We analyzed the change in strength of cohesin-independent links after auxin treatment by using APA at 100kb resolution. APA analysis clearly demonstrated that while cohesin-independent links (both intra and interchromosomal) were weakly present before auxin treatment, they were ~2-fold strengthened after auxin-treatment (Fig S5E). This result was robust to using either our manually curated lists (of 61 intra and 203 inter chromosomal links) or automatedly generated lists (all intrachromosomal pairs of the 47 superenhancers overlapping anchors in the 88-loop automated list from above and all interchromosomal pairs of the 47 superenhancers overlapping anchors in the 88-loop automated list from above) (Fig S5D).

We also analyzed induction of cohesin-independent links across an auxin treatment and withdrawal time course. This analysis was performed as above with the cohesin-associated loops and loop domains, but at 100kb resolution instead of 10kb resolution and for both our 61 intrachromosomal links and our 203 interchromosomal links. The opposite pattern of cohesin-associate loop formation was seen; APA scores for cohesin-independent links rapidly increased upon auxin treatment and rapidly dropped upon auxin withdrawal (Fig. 4D). Similar results were seen upon performing the time course APA at 100kb resolution using all intrachromosomal pairs of the 47 superenhancer overlapping anchors in the 88-loop automated list from above (Fig. S5B).

#### Simulations of extrusion and compartmentalization

Simulations were run for 200,000 timesteps with only Lennard-Jones intermonomeric forces and then for 800,000 timesteps with 8 extrusion complexes. In the HOOMD-blue molecular dynamics package (Anderson et al., 2008; Glaser et al., 2015), temperature is set to 2.0 and gamma (viscosity) is set to 0.02. Contact maps and globules are shown from the final frame of simulation. In simulations of the auxin-treated condition, the final 800,000 timesteps were simulated without extrusion. All other parameters are as described in (Sanborn et al.).

CTCF and cohesin binding strengths were determined by integrating a Gaussian fit to ChIP-Seq data around every CTCF motif. Simulated extrusion binding strengths were determined by taking the geometric mean of the CTCF and cohesin binding strengths and renormalizing to a binding probability, as described in (Sanborn et al., 2015).

Each monomer was assigned to either an “A” or a “B” type. Lennard-Jones forces between different-type monomers was set to 98% the strength of LJ forces between same-type monomers. Because compartment transitions can only be defined in Hi-C maps at coarse resolutions (25kb and above), the compartment transition of each simulation replicate was varied randomly within 30kb (30 monomers) of defined transition points, which were set based on treated Hi-C maps.

#### Assessment of changes in transcription after cohesin loss

To look for signs of ectopic activation, we examined the 14,853 genes that were not expressed (RPKM<0.5) in untreated cells. We identified 2,145 genes that were significantly (adjusted p<0.05) changed by DESeq2. Of these genes, 1% (216) were ectopically activated after treatment (p<0.05, >30% change in RPKM, RPKM>0.5 in treated cells). In addition, 7% of these genes (1063) exhibited “leaky” transcription in treated cells: a larger PRO-Seq signal (p<0.05, >1.3 fold change difference) that fell short of the threshold for an expressed gene (i.e., RPKM was still below 0.5). 1.4% of these genes were significantly downregulated (>1.3-fold change), but it is unclear what reductions in expression at such low levels of expression mean biologically.

We next looked for changes in the 12,222 genes that were expressed (RPKM>0.5) in untreated cells (Fig. 6B). We identified 4,196 genes that were significantly changed (adjusted p<0.05) changed by DESeq2. Here again, most genes (87%, 10,615) exhibited similar levels of transcription after cohesin degradation (RPKM changed by less than 30%). The remaining genes (13%, 1607) showed a larger transcriptional effect (p<0.05, >30% change in RPKM). Stronger effects were seen, but less frequently: 64 genes (0.5%) showed a 2-fold change, and 2 genes showed a 5-fold change (Fig. 6B).

We identified 49 genes that were 1.75-fold downregulated with p<0.05 after auxin treatment. We noticed that many of the genes that were downregulated (by >1.75-fold) were located within 500kb of superenhancers (23 of 49, 4.8-fold enrichment compared to randomly shuffling the positions of the TSS of the 49 genes across the genome, Fig. 6C,D). Of these genes, 29% (14 of 49) were located with 500kb of one of the top 100 superenhancers (8.5-fold enrichment compared to randomly shuffling the positions of the TSS of the 49 genes across the genome). The overall distribution of distance to the nearest superenhancer was shifted significantly closer compared to randomly selected genes (Fig. 6C). Strikingly, these superenhancers were often located at the anchors of the cohesin-independent links seen in treated cells (8 of 19, a 13.7-fold enrichment).

To rule out the possibility that changes in gene expression were due to the auxin hormone itself, we performed PRO-Seq on HCT-116-CMV-OsTIR1 cells (HCT-116 cells with OsTIR1 at the *AAVS1<* locus but no mAID tag on any protein) before and after auxin treatment. Only 105 genes were detected as significantly different, and only 56 genes were detected as significantly different with at least a 1.3-fold change. This indicates that our results are not confounded by the auxin hormone itself.

To rule out the possibility that tagging RAD21 itself led to significant transcriptional consequences, we compared our auxin-treated PRO-Seq data to a control of untreated HCT-116-CMV-OsTIR1 cells. The following paragraphs are the analyses from above except with the numbers from the CMV-OsTIR1 control. Analogous plots to those shown in Fig. 6B and 6C for the CMV-OsTIR1 control are shown in Fig. S7C-D.

**Figure S6:**
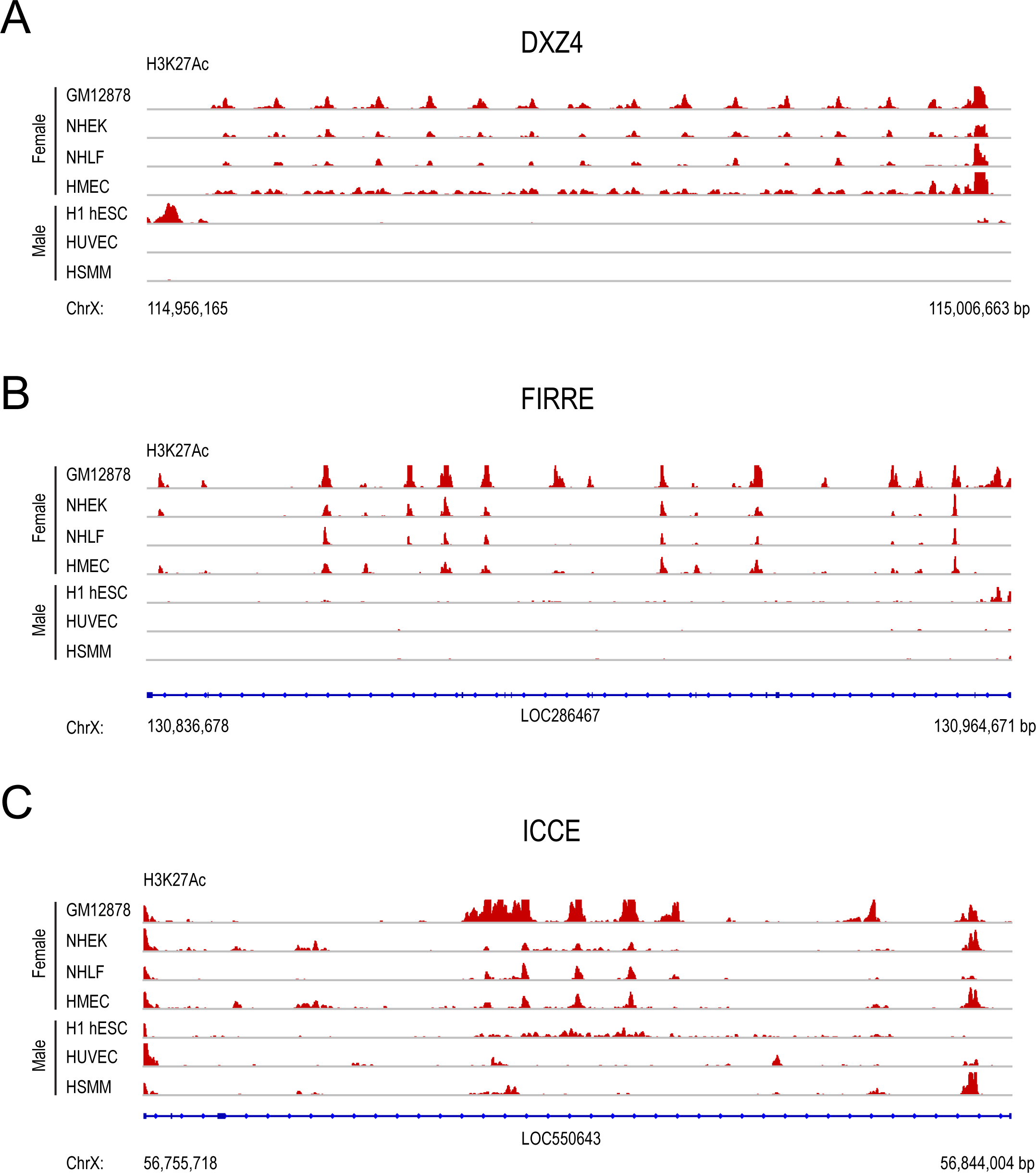
Superloop anchors on the inactive X chromosome are enriched for H3K27Ac in an allele-specific manner, Related to Figure 4. (A,B,C): DXZ4 (A), FIRRE (B) and ICCE (C), the three most prominent superloop anchors on the inactive X chromosome (Rao et al., 2014; Darrow et al., 2016) are enriched for H3K27Ac in female cell lines (GM12878, NHEK, NHLF, HMEC) but not male cell lines (H1-hESC, HUVEC, HSMM). All H3K27Ac tracks shown were generated by ENCODE (ENCODE Consortium, 2012) and are shown with a common maximum enrichment of 50.

**Figure S7:**
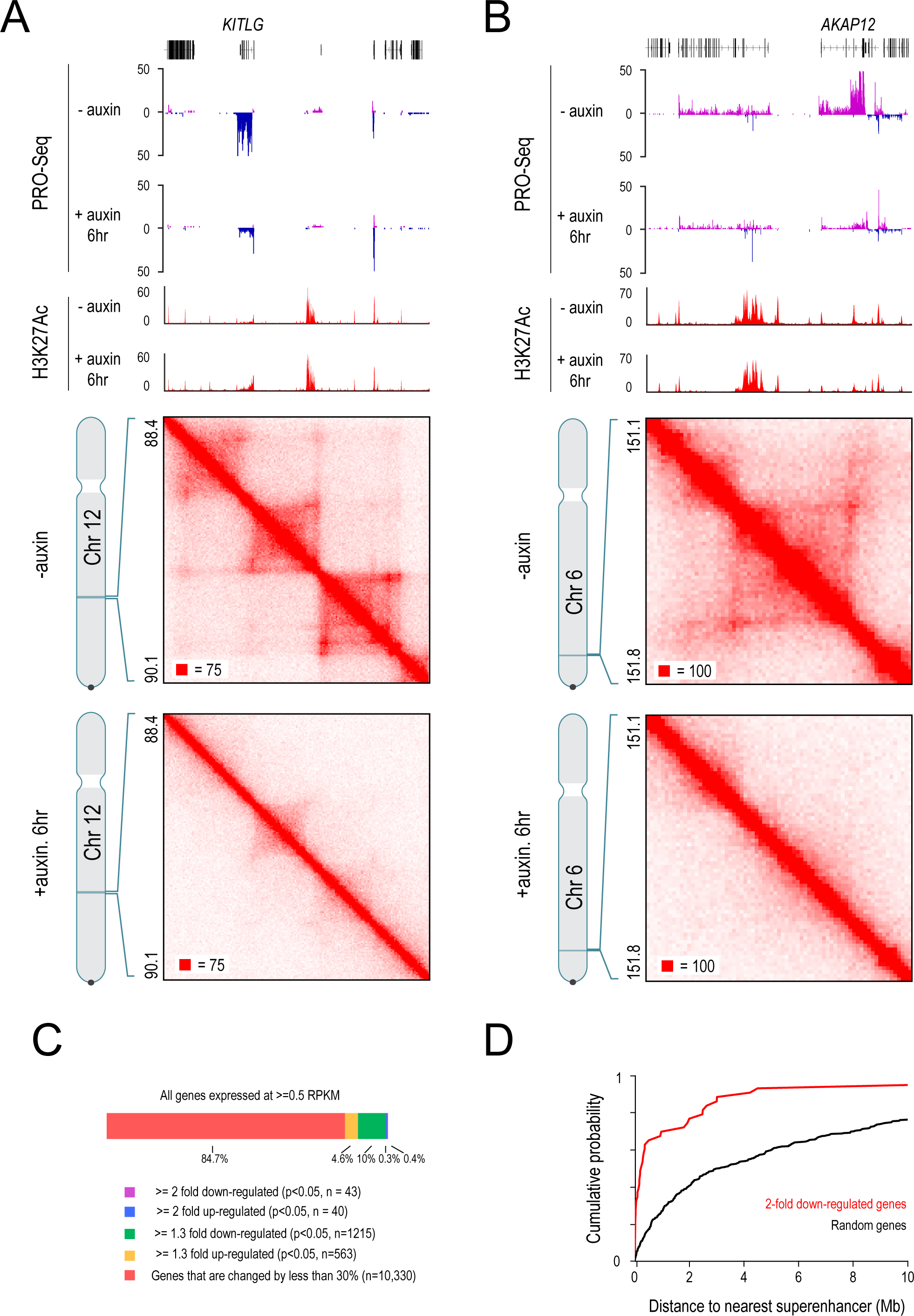
Examples of downregulation of genes nearby superenhancers after cohesin loss, Related to Figure 6. (A) An example of a strongly down-regulated gene near a superenhancer. In untreated cells, *KITLG* is contained within a loop domain with a strong superenhancer. Upon auxin treatment, the spanning loops are lost and *KITLG* expression is 2.73-fold down-regulated. The superenhancer near *KITLG* forms strong links to other superenhancers intrachromosomally and interchromosomally after auxin treatment. (B) An example of a strongly down-regulated gene near a superenhancer. In untreated cells, *AKAP12* is contained inside a loop domain with a strong superenhancer. Upon auxin treatment, the spanning loop is lost and *AKAP12* expression is 3.3-fold down-regulated. The superenhancer near *AKAP12* forms strong links to other superenhancers intrachromosomally and interchromosomally after auxin treatment. (C) Genes that are expressed in untreated cells rarely undergo substantial changes in expression level after cohesin loss even when compared to untagged HCT-116 CMV-OsTIR1 cells. (D) Cumulative probability distributions of distances to the nearest superenhancer for 2-fold down-regulated genes between untreated HCT-116 CMV-OsTIR1 cells and treated RAD21-mAC cells (red) versus random genes (black).

To look for signs of ectopic activation, we examined the 14,884 genes that were not expressed (RPKM<0.5) in untreated cells. We identified 2,284 genes that were significantly (adjusted p<0.05) changed by DESeq2. Of these genes, 1% (255) were ectopically activated after treatment (p<0.05, >30% change in RPKM, RPKM>0.5 in treated cells). In addition, 7% of these genes (1179) exhibited “leaky” transcription in treated cells: a larger PRO-Seq signal (p<0.05, >1.3 fold change difference) that fell short of the threshold for an expressed gene (i.e., RPKM was still below 0.5). 1.8% of these genes were strongly downregulated (>1.3-fold change), but it is unclear what reductions in expression at such low levels of expression mean biologically.

We next looked for changes in the 12,191 genes that were expressed (RPKM>0.5) in untreated cells (Fig. 5B). We identified 4,251 genes that were significantly changed (adjusted p<0.05) changed by DESeq2. Here again, most genes (85%, 10,330) exhibited similar levels of transcription after cohesin degradation (RPKM changed by less than 30%). The remaining genes (15%, 1861) showed a larger transcriptional effect (p<0.05, >30% change in RPKM). Stronger effects were seen, but less frequently: 86 genes (1%) showed a 2-fold change, and 3 genes showed a 5-fold change (Fig. S7C).

We identified 43 genes that were 2-fold downregulated with p<0.05 after auxin treatment. We noticed that many of the genes that were downregulated (by >2-fold) were located within 500kb of superenhancers (28 of 43). Of these genes, 49% (21 of 43) were located with 500kb of one of the top 100 superenhancers. The overall distribution of distance to the nearest superenhancer was shifted significantly closer compared to randomly selected genes (Fig. S7D).

## DATA AND SOFTWARE AVAILABILITY

All datasets reported in this paper are available at the Gene Expression Omnibus (GEO), series accession number GSEXXXX.

## ADDITIONAL RESOURCES

KEY RESOURCES TABLE

**KEY RESOURCES TABLE**
| REAGENT or RESOURCE | SOURCE | IDENTIFIER |
| --- | --- | --- |
| Antibodies |  |  |
| Rabbit Polyclonal anti-SMC1 | Bethyl Laboratories | A300-055A |
| Rabbit Polyclonal anti-RAD21 | Abcam | ab992 |
| Rabbit Monoclonal anti-CTCF | Cell Signaling | 3418 |
| Rabbit Polyclonal anti-H3K27Ac | Abcam | ab4729 |
| Rabbit Monoclonal anti-H3K4me3 | Millipore | 04-745 |
| Rabbit Polyclonal anti-H3K4me1 | Abcam | ab8895 |
| Rabbit Polyclonal anti-H3K36me3 | Abcam | ab9050 |
| Rabbit Polyclonal anti-H3K27me3 | Millipore | 07-449 |
| Rabbit Polyclonal anti-H3K9me3 | Abcam | ab8898 |
| Rabbit Polyclonal anti-H4K16Ac | Millipore | 07-329 |
| Rabbit Polyclonal anti-H4K20me3 | Abcam | ab9053 |
| Rabbit Polyclonal anti-H3K79me2 | Abcam | ab3594 |
| Rabbit Polyclonal anti-H2A.Z | Abcam | ab4174 |
| Chemicals, Peptides, and Recombinant Proteins |  |  |
| 3-indole acetic acid | Sigma Aldrich | I3750-5G-A |
| Deposited Data |  |  |
| Raw and analyzed data | This paper | GSEXXX |
| ENCODE ChIP-Seq peak calls for DNA binding proteins and histone modifications in HCT-116 | ENCODE;<br>encodeproject.org | https://www.encodeproject.org/search/?searchTerm=HCT-116&type=Experiment&assay_slims=DNA+binding |
| Stitched superenhancers and enhancers in HCT-116 | Hnisz et al., 2013 | http://www.sciencedirect.com/science/article/pii/S0092867413012270; Table S2 |
| Hi-C data from Seitan et al., 2013 | Seitan et al., 2013 | GEO GSE48763 |
| Hi-C data from Sofueva et al., 2013 | Sofueva et al., 2013 | GEO GSE49017 |
| Hi-C data from Zuin et al., 2014 | Zuin et al., 2014 | GEO GSE44267 |
| Experimental Models: Cell Lines |  |  |
| HCT-116 CMV-OsTIR1 | Laboratory of M. Kanemaki; Natsume, et al., 2016 | N/A |
| HCT-116 RAD21-mAiD-mClover | Laboratory of M. Kanemaki; Natsume, et al., 2016 | N/A |
| Software and Algorithms |  |  |
| Juicer | Durand et al., 2016 | http://aidenlab.org/documentation.html |
| Juicebox | Durand et al., 2016 | http://aidenlab.org/documentation.html |
| BWA-MEM | Li and Durbin, 2010 | http://bio-bwa.sourceforge.net/ |
| MACS2.0 | Liu, 2014 | https://github.com/taoliu/MACS |
| Bowtie | Langmead and Salzberg, 2012 | bowtie2 v2.1.0<br>http://bowtie-bio.sourceforge.net/index.shtml |
| Samtools | Li et al., 2009 | samtools v.0.1.19<br>https://github.com/samtools/samtools |
| Picard Tools |  | http://picard.sourceforge.net<br>https://broadinstitute.github.io/picard/ |
| bwtool | Pohl and Beato, 2014 | https://github.com/CRG-Barcelona/bwtool/wiki |
| DESeq2 | Love et al., 2014 | http://bioconductor.org/packages/release/bioc/html/DESeq2.html |
| Other |  |  |

